# SUN-domain proteins of the malaria parasite *Plasmodium falciparum* are essential for proper nuclear division and DNA repair

**DOI:** 10.1101/2024.04.23.590856

**Authors:** Sofiya Kandelis-Shalev, Manish Goyal, Tal Elam, Shany Assaraf, Noa Dahan, Omer Farchi, Eduard Berenshtein, Ron Dzikowski

## Abstract

The protozoan parasite *Plasmodium falciparum*, which is responsible for the deadliest form of human malaria, accounts for over half a million deaths a year. These parasites proliferate in human red blood cells by consecutive rounds of closed mitoses called schizogony. Their virulence is attributed to their ability to modify the infected red cells to adhere to the vascular endothelium and to evade immunity through antigenic switches. Spatial dynamics at the nuclear periphery were associated with regulation of processes that enable the parasites to establish long-term infection. However, our knowledge of components of the nuclear envelope (NE) in *Plasmodium* remains limited. One of the major protein complexes at the NE is the LINC complex that forms a connecting bridge between the cytoplasm and the nucleus through the interaction of SUN and KASH domain proteins. Here we have identified two SUN-domain proteins as possible components of the LINC complex of *P. falciparum* and show that their proper expression is essential for the parasite’s proliferation in human red blood cells and that their depletion leads to the formation of membranous whorls and morphological changes of the NE. In addition, their differential expression highlight different functions at the nuclear periphery as PfSUN2 is specifically associated with heterochromatin, while PfSUN1 expression is essential for activation of the DNA damage response. Our data provide indications for the involvement of the LINC complex in crucial biological processes in the intraerythrocytic development cycle of malaria parasites.

## Introduction

Malaria caused by protozoan parasites of the genus *Plasmodium* remains one the leading causes of morbidity and mortality worldwide. These parasites are estimated to infect over 200 million people each year resulting in over 600,000 documented deaths (1). *Plasmodium falciparum* is responsible for the deadliest form of human malaria that accounts for over 90% of malaria deaths, primarily of young children and pregnant women. The lack of an effective vaccine as well as the ability of the parasite to develop resistance to effectively all anti-malarial drugs, keeps malaria as a major health and economic burden in many endemic countries.

*P. falciparum* has a complex life cycle, during which it undergoes unique replication cycles and morphological changes to adapt to different hosts and changing environments. These changes are mediated by tight regulation of cell cycle-dependent gene expression patterns (2) (3). Like any other eukaryote, the parasite’s ability to proliferate and thrive in its human host depends on precise orchestration of biological processes and molecular exchange between the cell’s nucleus and the cytoplasm, which are separated by the nuclear envelope (NE). Evidently, the NE isn’t only a passive nuclear boundary, it was shown to be involved in controlling nuclear and cellular processes and plays critical roles in transferring signals from the cytoplasm and the extra-cellular environment into the nucleus (4–7).

One of the major protein complexes that connects nuclear and cytoplasmic processes is the Linker of Nucleoskeleton and Cytoskeleton (LINC) protein complex (8–11). The LINC complex spans the Nuclear Envelope (NE) across the fluid lumen and forms a direct mechanical connection between the nucleus and the cytoplasm. LINC complexes are comprised of SUN domain proteins, residents of the Inner Nuclear Membrane (INM), that interact with KASH domain proteins (Nesprins) that are embedded in the Outer Nuclear Membrane (ONM). In higher eukaryotes, the LINC complex was demonstrated to form a physical contact between components of the cytoskeleton and the nuclear lamina, and was implicated in determining nuclear positioning, migration and orientation as well as influencing cell division (12). In addition, the LINC complex was implicated in DNA repair (13–15), and in chromosome organization by forming a mechanical link between the cytoskeleton and DNA (16–19). Furthermore, in addition to providing structural integrity for the nucleus, the LINC complex allows for the transfer of mechanical force into the nucleus resulting in mechanotransduction of signals from the cell’s extracellular environment to the nucleus (8, 20).

Very little is known about components of the NE in *P. falciparum.* These parasites lack conventional lamins, and thus, factors that constitute the nuclear scaffold under the INM remain unknown. In addition, only a few nucleoporins have been characterized, despite the functional conservation of the Nuclear Pore Complex throughout eukaryotic evolution. Similar to other components of the NE, little is known about constituents of the LINC complex in *Plasmodium* or any other apicomplexan parasites. Recent studies have revealed the presence of an alternative LINC-like complex in *P. berghei*, comprising SUN1 and the newly identified protein ALLC1, which serves a functionally analogous role to KASH-domain proteins (21, 22). The SUN1-ALLC1 complex has been implicated in the organization of the microtubule organizing center (MTOC) and the coordination of mitotic spindle formation, highlighting its critical role in male gametogenesis and malaria transmission.

Here we identified and characterized two *P. falciparum* SUN-domain proteins, which we called PfSUN1 and PfSUN2. We found that these NE proteins play a role in parasite proliferation during the intra-erythrocytic development cycle (IDC). Depletion of expression of both PfSUN1 and PfSUN2 results in increased cellular deformability during schizogony and the appearance of membranous whorls that bud from the NE at earlier stages. Interestingly, we found that PfSUN2 appears to be strongly associated with heterochromatin at the nuclear periphery. Finally, our data indicate that PfSUN1 depletion interferes with the parasite’s ability to activate the DNA damage response. Our data provide the first functional roles of possible components of the LINC complex in *P. falciparum* biology.

## Results

### Putative SUN domain proteins in *Plasmodium falciparum*

We performed a BLAST search in the *Plasmodium* database (https://plasmodb.org/plasmo/app) (23) using the amino acid sequences of the human orthologues and identified two putative proteins that contain SUN domains, which we named as PfSUN1 (PF3D7_1215100) and PfSUN2 (PF3D7_1439300; **Fig. S1**). We found that the SUN domain of PfSUN1 displayed higher similarity and structural conservation with that of the human HsSUN2 & 5 while the SUN-domain of PfSUN2 share higher similarity with that of *Arabidopsis thaliana* (**Fig. S1A, B, D & E**). Both proteins are conserved among *Plasmodium* species (**Fig. S1 C & F**). Multiple sequence alignments of PfSUN1 and PfSUN2 with different SUN domains from evolutionarily distinct organisms indicated that the SUN domains could be separated into two main clades (**Fig. S1G** left). Interestingly, while PfSUN1 belongs to the same clade as the canonical C’ SUN domain family, PfSUN2 appears to resemble mid SUN-domain proteins characterized primarily in plants (**Fig. S1G** right). In both cases, *P. falciparum* SUN-domain proteins contain the structural features of their respective protein clade i.e. trans-membrane (TM) domains, coiled-coiled domains (CC), and the SUN domain. This conservation suggests that Plasmodial putative SUN domain proteins may function as components of the LINC complex, similar to SUN domain proteins in other organisms.

### PfSUN1 is a nuclear envelope protein

Analysis of the available transcriptomic data in (https://plasmodb.org/plasmo/app) indicated that both SUN-domain proteins are transcribed throughout the parasite’s life cycle with some stage-specific variations in their transcript levels (**Fig. S2**). To begin characterizing *P. falciparum* putative SUN-domain proteins we initially expressed PfSUN1-GFP ectopically (Fig. S3A, B) and followed its cellular localization during the parasite’s intra-erythrocytic development (IDC). We found that PfSUN1-GFP surrounds the DNA staining at the nuclear periphery throughout the IDC (**Fig. S3B**), suggesting possible localization with the nuclear envelope (NE). Interestingly, in some nuclei PfSUN1-GFP is concentrated at distinct foci at the nuclear periphery. To validate its association with the NE we performed immune-EM and found that PfSUN1-GFP is indeed located at the NE (**Fig. S1C**). We performed dSTORM imaging to test its association with the NPC using the nucleoporin PfSec13 as a marker (24). To this end, parasites were transfected with two different episomes expressing either PfSUN1 fused with GFP (PfSUN1-GFP) and PfSEC13 fused with HALO-tag (PfSec13-Halo). we found that both PfSUN1 and PfSec13 surround the nuclear periphery during the IDC (**Fig. 1A**). Interestingly, while PfSec13 localized to the distinct foci of the nuclear pore complexes (NPC) as previously demonstrated, PfSUN1 appears to be surrounding the nucleus in a more continuous pattern, which also contains foci of stronger signal that are often adjacent to the NPC. We have previously observed changes in the directionality of the NPC as mitosis progress that provided evidence for cytoplasmic influence on NPC localization during the transition from a multinucleated syncytium to multiple autonomous cells (25). Interestingly, we observed pronounced accumulation of PfSUN1 in late stage schizonts at the same nuclear pole with the NPC clusters (**Fig. 1B**). The nuclear periphery of *P. falciparum* includes regions of condensed heterochromatin marked by the histone modification H3K9me3. This histone modification appears to be an epigenetic marker specifically devoted to gene families that undergo clonally variant transcription that localize to the nuclear periphery (3, 26). Further dSTORM imaging indicated that while it surrounds the nucleus, PfSUN1-GFP does not co-localize with either the heterochromatic marker H3K9me3 or the euchromatin marker H3K9ac (**Fig. 1C & D**).

**Figure 1.**
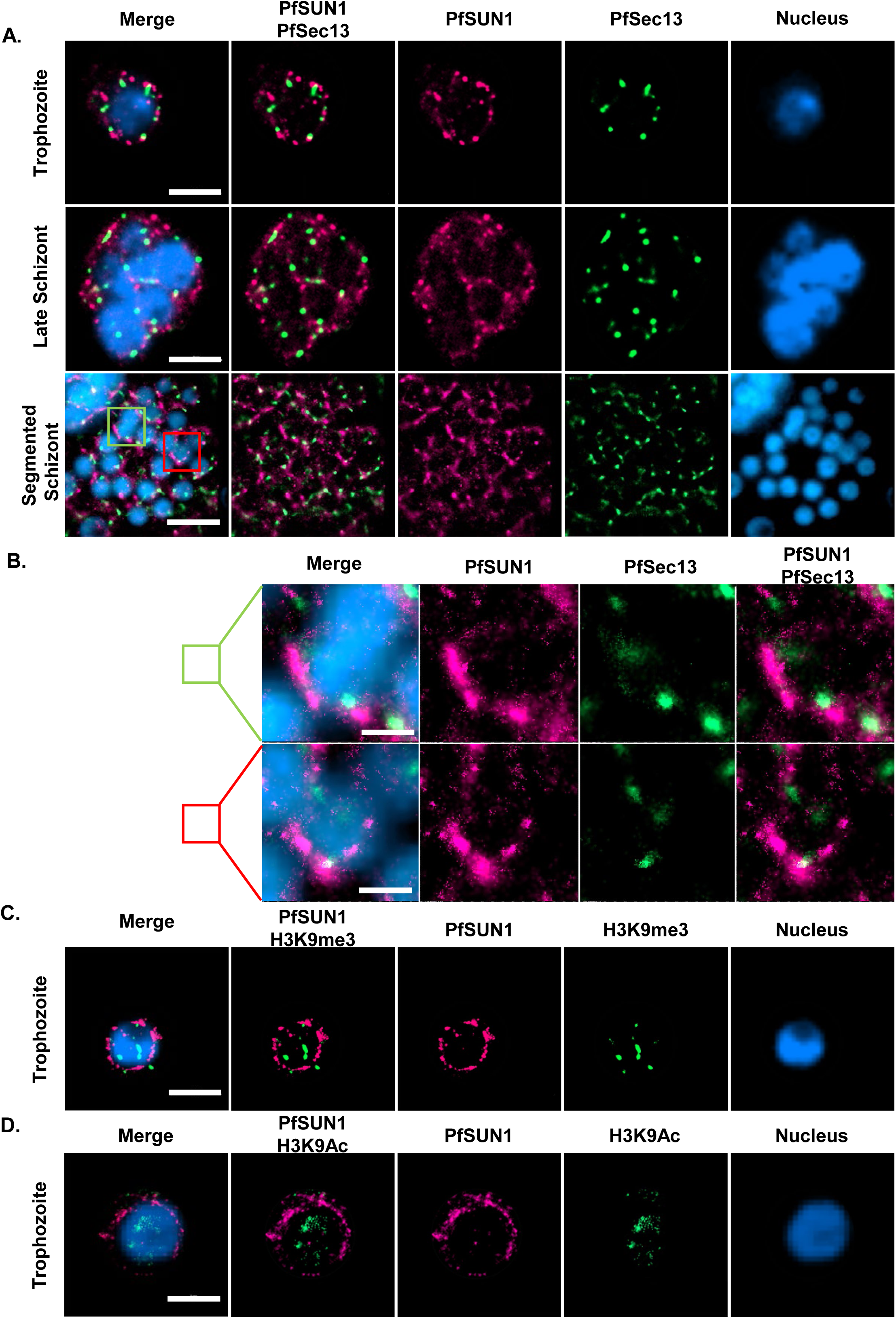
Ectopic PfSUN1 is localized to the nuclear periphery and displays polar accumulation during schizogony. **A.** Dual color dSTORM imaging analysis of PfSUN1-GFP (magenta), with PfSEC13-Halo (green) during *P. falciparum* IDC showing their distribution at the nuclear periphery during IDC. Nuclei were stained with YOYO1® dye (blue). Scale bar: 2 µm. **B.** High magnification of merozoites from the segmented schizont labeled by green or magenta rectangle show polar distribution of the NPC and PfSUN1-GFP. Scale bar: 0.5 µm. **C.** dSTORM imaging of PfSUN1 association with heterochromatin and euchromatin histone markers. Upper panel shows PfSUN1-GFP (magenta) localization with the heterochromatin mark H3K9me3 (yellow), and the lower panel shows localization with the euchromatin mark H3K9Ac (green). Scale bar: 2 µm.

To determine which of the PfSUN1 domains is essential for its nuclear positioning we performed deletion analysis of the coiled-coiled and the SUN domains and imaged parasites expressing different truncated versions of PfSUN1 fused with a *myc* epitope tag (**Fig. 2**). We found that deletions of the coiled-coiled domain as well as deletion of both the coiled-coil and the SUN domain abolished the positioning of PfSUN1 at the nuclear envelope, even though the trans-membrane domain was not deleted. These data suggest that the coiled-coil domain is essential for proper positioning of PfSUN1 at the NE.

**Figure 2.**
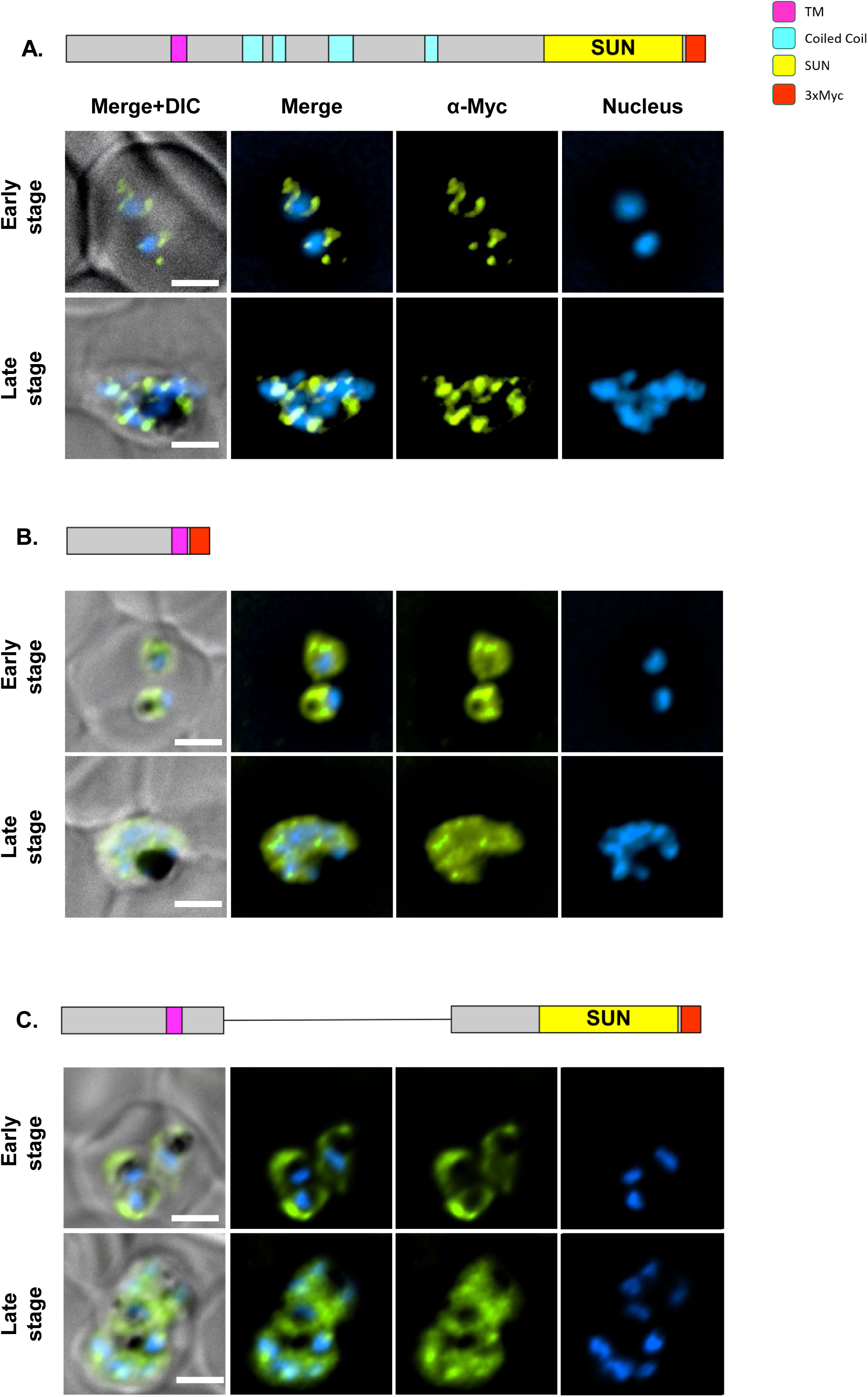
Deletion of PfSUN1 coiled coil motifs results in the loss of nuclear envelope localization. Immunofluorescence imaging of parasite that ectopically express different deletion mutants of PfSUN1-myc in NF54 parasites. **A.** Full length PfSUN1-myc. **B.** PfSUN1 1-190-myc (expressing the N terminal 190 aa of PfSUN1). **C.** PfSUN1Δ250-600-myc (coiled coil domains deleted). Anti-myc antibody labels PfSUN1 mutants (green), nuclei were stained with DAPI (blue). Predicted domain architecture: SUN domain (yellow), transmembrane domain (magenta), coiled coil domains (cyan), myc tag (red). Scale bars: 2μm. DIC, Differential interference contrast.

To better understand the role of the two putative SUN-domain proteins in *P. falciparum* we used the pSLI system (27) to create transgenic parasite lines in which PfSUN1 or PfSUN2 are endogenously tagged with an HA epitope and fused to the *glmS* ribozyme that allows one to perform conditional knock-down by adding glucosamine (GlcN) to the culture media. These lines were termed PfSUN1-HA-*glmS* and PfSUN2-HA-*glmS,* respectively. Following the isolation of a clonal transgenic population and confirming the correct integration into the genome (**Fig. S4 & Fig. S5**) we imaged their cellular localization to confirm that the endogenously tagged PfSUN1 localized to the nuclear periphery. We initially performed IFA using anti BiP as a marker for the ER that in *P. falciparum* appears to be surrounding the nuclear staining. Indeed, we found that PfSUN1 localizes to the nuclear periphery with a no complete overlap with the ER. To gain more accurate imaging of PfSUN1 and overcome the diffraction limits of light microscopy, we used dSTORM imaging to confirm that the endogenously tagged PfSUN1 localized to the nuclear periphery (**Fig. 3A & B**) and was not associated with the heterochromatin marker H3K9me3 (**Fig. 3C**) as observed in its ectopic expression. Similar results were obtained for the endogenously tagged PfSUN2 (**Fig. 4**). PfSUN2 is expressed at the nuclear periphery during the IDC, demonstrating regions of stronger signal where the protein appears to be accumulated (**Fig 4B & C**). Interestingly, our dSTORM imaging data suggest that the foci of PfSUN2 are associated with foci of heterochromatin (marked by H3K9me3 antibody) at the nuclear periphery (**Fig. 4C**). Using 5 mM GlcN we were able to downregulate the expression of PfSUN1 and PfSUN2 (**Fig. 5A & B)** and show that this knock-down decreased the proliferation rate of the cultured parasite population (**Fig. S6 A & B)**. We normalize the basal growth rate of each line and the effect of GlcN on growth, by calculating the ratio of parasitemia of each line with or without GlcN, and show that knock-down of both PfSUN1 and PfSUN2 causes a significant growth delay (**Fig. 5C**), associated with decrease in the number of nuclei per schizont and an increase in the number of parasites with aberrant nuclei segregation (Fig. 5D & Fig. S10). Altogether, these data demonstrate that PfSUN1 and PfSUN2 are NE proteins, which are essential for proper proliferation of *P. falciparum*.

**Figure 3.**
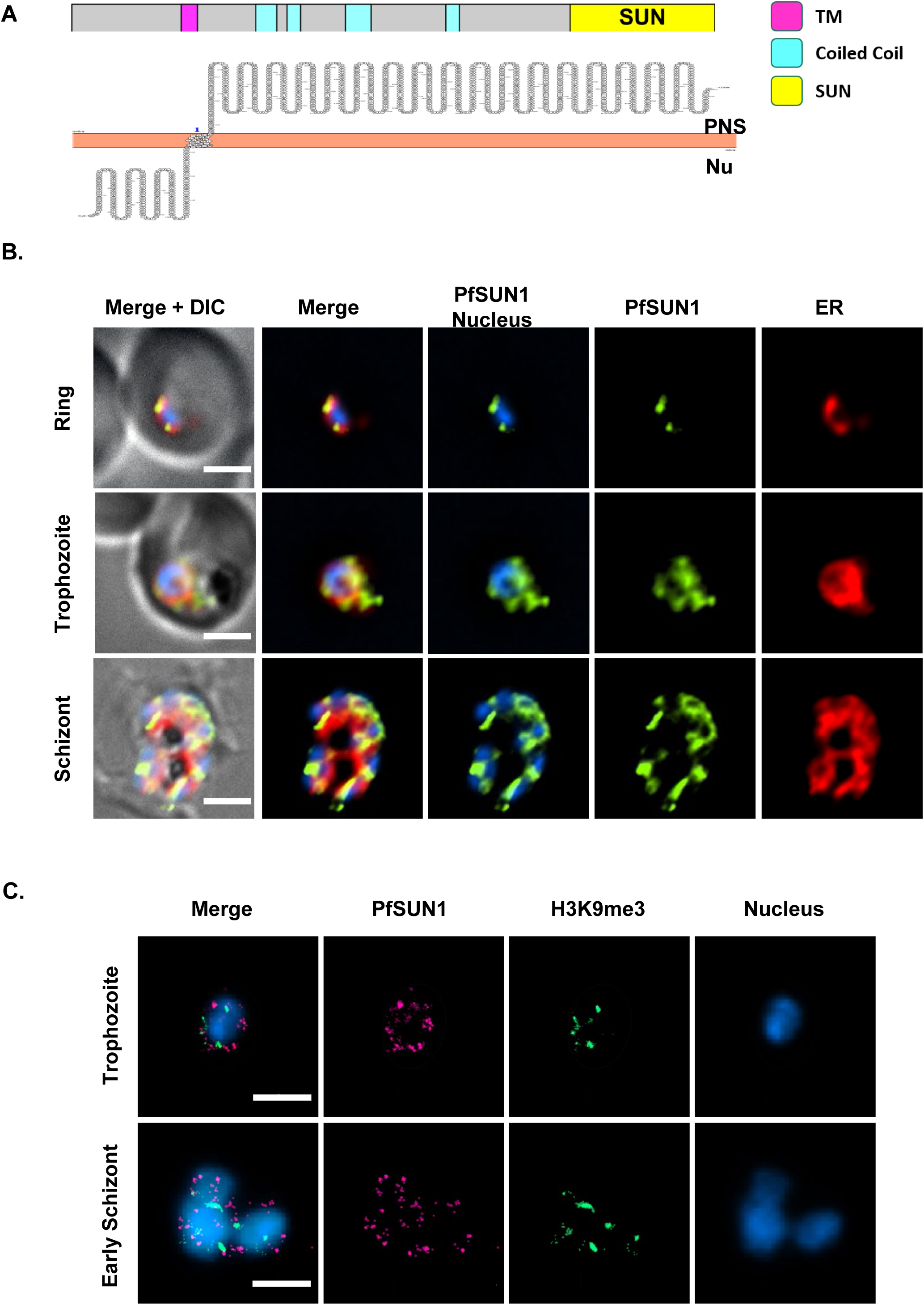
Endogenous PfSUN1 is not associated with heterochromatin at the nuclear periphery. **A.** Schematic representation of domain architecture of PfSUN1 (SUN domain; yellow, transmembrane domain; magenta, coiled coil domains; cyan), membrane topology prediction was analyzed using Protter (54) (PNS; Perinuclear Space. Nu; Nucleoplasm). **B.** IFA imaging demonstrating the expression of endogenous PfSUN1-HA during different stages of IDC in PfSUN1-HA-*glmS* transgenic parasites. PfSUN1-HA (green), ER marker (BiP, red), nuclei were stained with DAPI (blue). Scale bar: 2 µm. **C.** dSORTM imaging of endogenous HA-tagged PfSUN1 (magenta) along with heterochromatin histone mark H3K9me3 (green). Nuclei were stained with YOYO1®stain (blue). Scale bar: 2µm.

**Figure 4.**
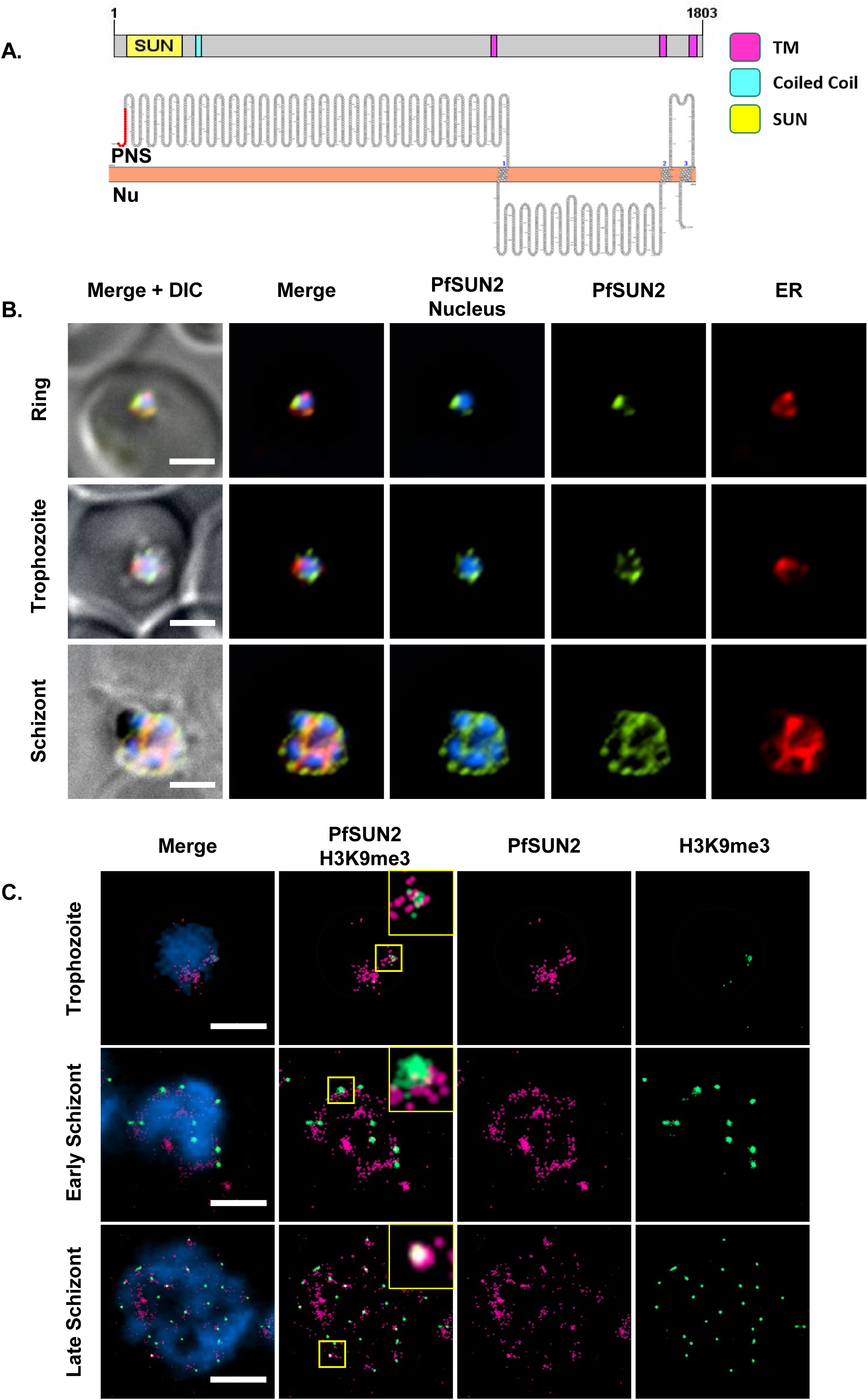
PfSUN2 is a nuclear envelope protein associated with heterochromatin. **A.** Schematic representation of domain architecture of PfSUN2 (SUN domain (yellow); transmembrane domain (magenta); coiled coil domains (cyan). Membrane topology prediction was analyzed using Protter (54) (PNS, Perinuclear Space; Nu, Nucleoplasm). **B.** IFA imaging demonstrating the distribution of endogenous PfSUN2-HA during different stages of IDC in PfSUN2-HA-*glmS* transgenic parasites. PfSUN2-HA (green), ER marker (BiP, red), nuclei were stained with DAPI (blue). Scale bar: 2µm. **C.** dSTORM imaging of endogenous HA-tagged PfSUN2 (magenta) along with heterochromatin histone mark H3K9me3 (green), nuclei were stained with YOYO1® (blue). Scale bar: 2µm. insert: higher magnification of marked foci.

**Figure 5.**
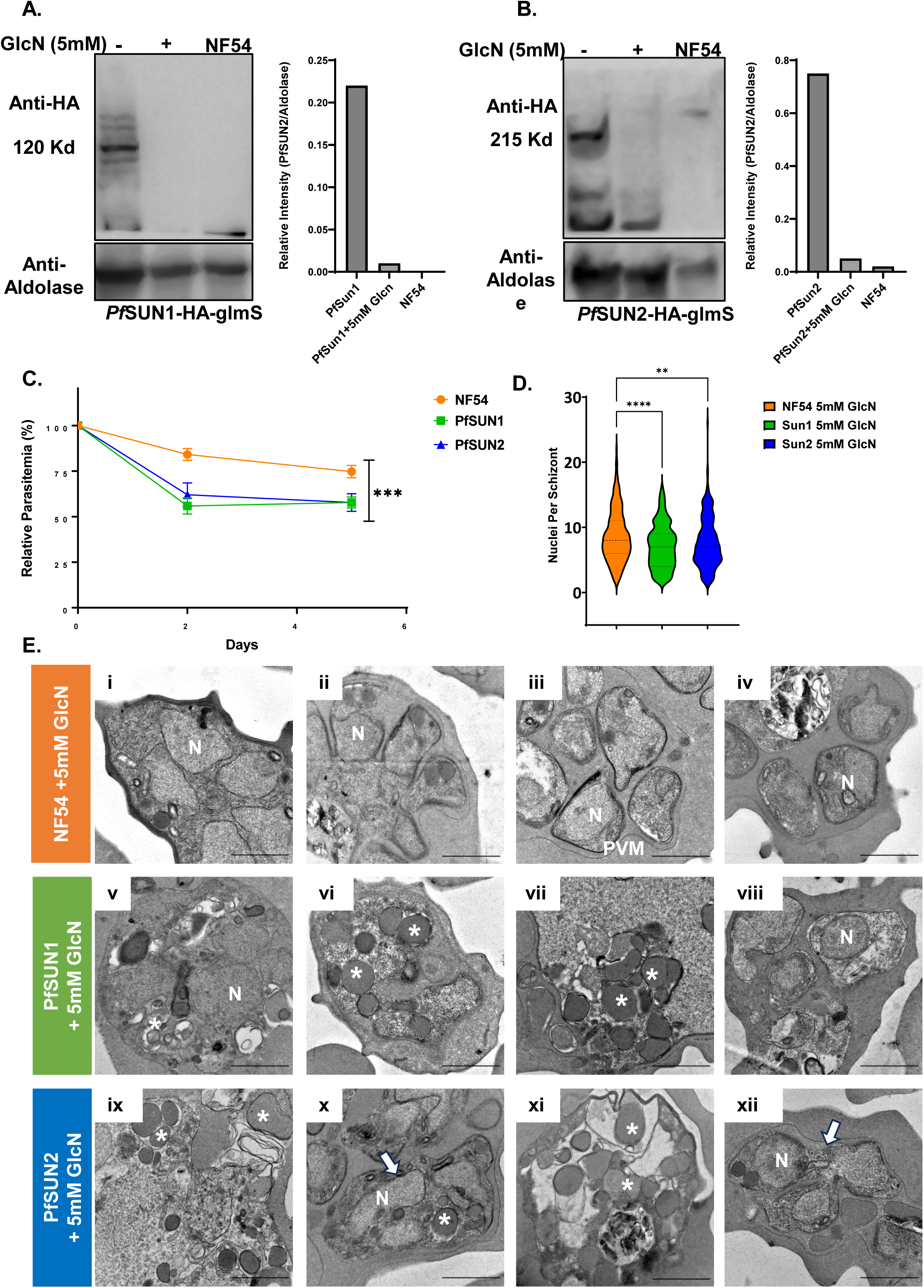
PfSUN1 and PfSUN2 expression is essential for the proper proliferation of blood-stages parasites. **A.** Inducible knock-down of endogenous PfSUN1. WB analysis demonstrating inducible knock-down by growing PfSUN1-HA-*glmS* parasites in the presence or absence of 5 mM GlcN for 72h. PfSUN1 was detected using an anti-HA antibody while an Anti-Aldolase antibody was used as a loading control. **B.** Inducible knock down of endogenous PfSUN2. WB analysis of PfSUN2-HA-*glmS* parasites grown in the presence or absence of 5 mM GlcN over a 72h time course. PfSUN2 was detected using anti-HA antibody, while anti-Aldolase antibody was used as a loading control. **C.** Inducible Knock-down of PfSUN1 & PfSUN2 cause significant growth delay. PfSUN1-HA-*glms*, PfSUN2-HA-*glms* or NF54 WT parasites were grown either in the presence or the absence of 5mM GlcN, with parasitemia determined daily by flow cytometry. Growth delays is expressed as the ratio of parasitemia at each time point between the GlcN treated and untreated parasites for each of the 3 lines. Experiments were performed in biological triplicates (n=3). Statistical significances between different stages were determined using student’s t-test (indicates *** P<0.001). **D.** Violin plots shows the distribution of nuclei per schizont, with the median (thick dashed line), and 25% and 75% quartile ranges indicated (thin dashed lines) for NF54, PfSUN1, and PfSUN2 lines treated with 5 mM GlcN. Data from Giemsa-stained blood smears of tightly synchronized parasites of each line used for EM, was obtained from 200–300 schizonts, and counted independently by two observers. Statistical significance was determined using pairwise comparisons; ** p < 0.01, **\*\*\*\*** p < 0.0001. **E.** Ultrastructure analysis using Transmission Electron Microscopy of schizonts grown in the presence of 5 mM GlcN. **i-.** NF54 wild type parasites showing normal morphology in pre-segmentation **(i)** mid segmentation **(ii)** parasitophorous vacuole (PVM) rupture **(iii-iv)**. PfSUN1-HA-*glmS* (**v-viii)** and PfSUN2-HA-*glmS* parasites (**ix-xii**) grown on GlcN for 96h to induce complete knockdown of PfSUN1 and PfSUN2. Representative micrographs showing abnormal morphology of pre- and post-segmented schizont with numerous membrane-bound vesicles in its cytoplasm, appearance of electron dense membrane bound structures, loss of internal structures and impaired segmentation. Asterisks indicate vacuolar structure and arrows point to unsegmented nuclei. scale bar: 1µm. N: Nucleus, PVM; Parasitophorous Vacuole.

### Depletion of PfSUN1 and PfSUN2 results in morphological alterations of the parasites

SUN domain proteins were shown to play an important role in maintaining nuclear architecture (28). We hypothesized that the reduction in growth rates of parasite populations in which PfSUN1 and PfSUN2 expression was knocked-down could imply that some of these parasites are unable to segregate and produce viable daughter merozoites. To better understand if this growth phenotype and its associated morphological changes observed by Giemsa stains (Fig. S10), we used Transmission Electron Microscopy (TEM) to evaluate the ultrastructural changes in the parasites’ morphology during replication. We found that while NF54 parasites appear to be able to divide their nuclei and segregate properly (**Fig. 5E**, upper panel), significant morphological deformations and loss of internal structures are observed during the progression of schizogony in many (but not all) of the parasites in which PfSUN1 (**Fig. 5E**, middle panel) and PfSUN2 (**Fig. 5E**, lower panel) are down-regulated. Large electron dense membrane bound vacuolar structures could be observed in these parasite populations, which may represent an apoptotic-like phenotype of parasites that will not be able to complete their intraerythrocytic development. These data indicate that SUN domain proteins in *P. falciparum* play a role in proper nuclei segregation and cellular division into daughter merozoites during schizogony.

As part of the LINC complex, SUN domain proteins that are embedded in the NE could play a role as a scaffold that holds the INM and ONM of the NE. We therefore investigated whether depletion of PfSUN1 and PfSUN2 would affect the morphology of the NE. To this end we used TEM on tightly synchronized trophozoites and were able to detect numerous deformations in the NE of parasites in which PfSUN2 expression is knocked down (**Fig. 6A & B**). However, the NE of the PFSUN1 knock-down parasite appeared to have normal morphology. In particular, we found that the average measured distance between the INM and the ONM was significantly wider in the PfSUN2 knock-down parasites. On the other hand, we found that PfSUN1 knock-down significantly reduced nuclei circularity, while PfSUN2 showed no significant impact (**Fig. 6C**). We also measured additional morphological parameters such as nuclei size, nuclei/cell ratio, food vacuole size and the ratio between the size of the food vacuole and the entire cell. In any of these parameters we were unable to detect significant change resulting from PfSUN1 and PfSUN2 knock-down (**Fig 6D-G**). Interestingly, in both PfSUN1 and PfSUN2 knock down parasites we observed duplication of the NE membranes (**Fig. 7A, v & ix, Fig. S8A & S9A** respectively). In addition, we observed the formation of concentric lamellar membranous whorls that involve both the INM and ONM, and in some parasites these whorls appeared to be budding from the NE (**Fig. 7A, vi & x, Fig. S8B & S9B** respectively), while they were also observed in the cytoplasm (**Fig. 7A, vii & xi, Fig. S8C & S9C** respectively) or the food vacuole (FV) (**Fig. 7A, viii & xii, Fig. S8D & S9D** respectively). Such membranous structures were also observed in NF54 parasites (Fig. 7A i-iv & Fig. S7), however, their abundance was more pronounced following PfSUN1 knockdown, with 40% of the parasites featuring them, as compared to 27% in the PfSUN2 knock down and only 14% of the NF54 parasites (**Fig. 7B**). In addition, we observed that the area of the membranous whorls in our thin sections varied between parasites (**Fig. 7C**).

**Figure 6.**
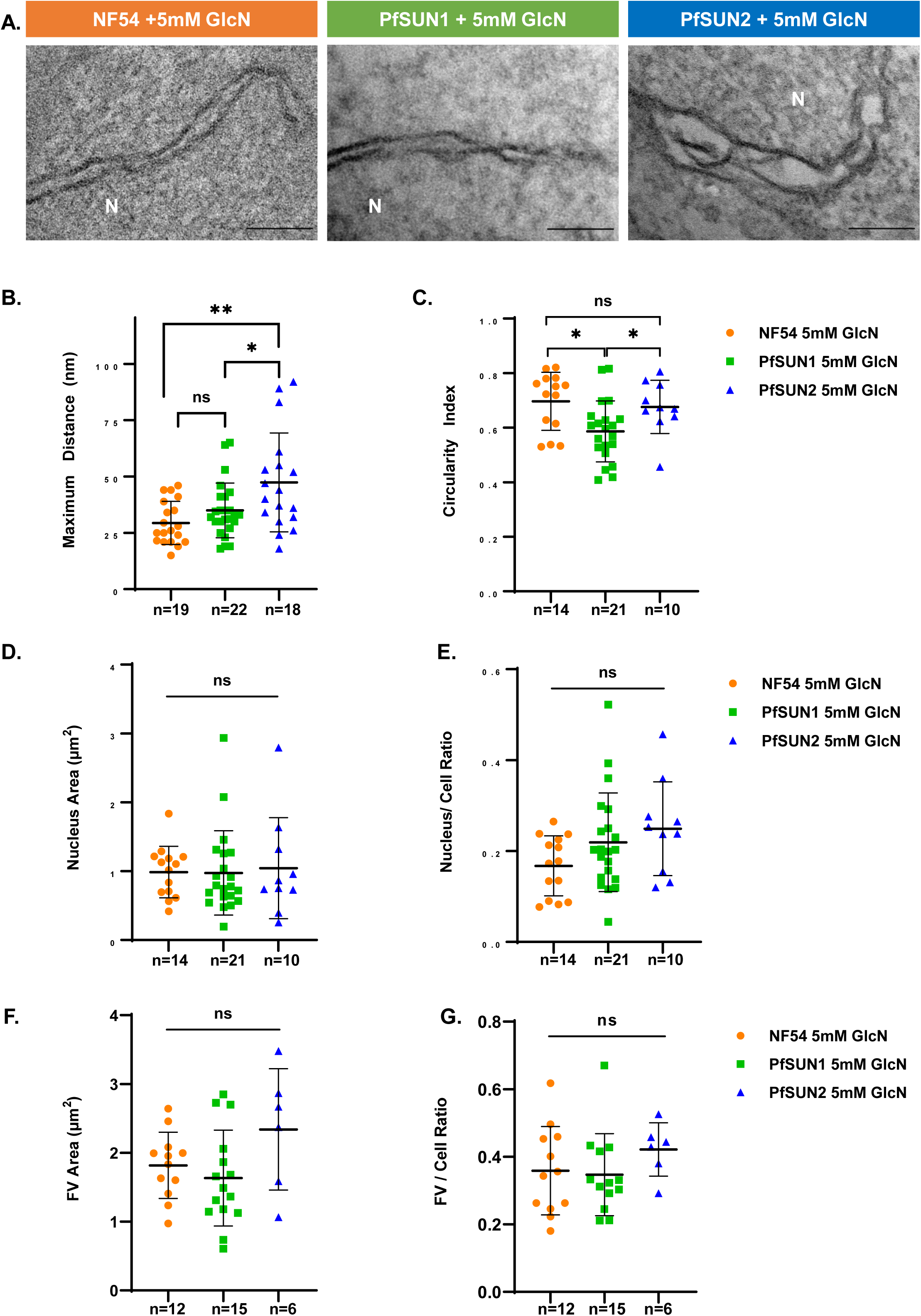
Loss of PfSUN1 and PfSUN2 expression impairs nuclear envelope homeostasis. Ultrastructural analysis of the nuclear and cellular morphology of late trophozoite of PfSUN1-HA-*glmS* and PfSUN2-HA-*glmS* parasites grown on 5 mM GlcN for 82h was examined using TEM. NF54 wild type parasites were grown in parallel on 5 mM GlcN for 82h as a control. **A.** Representative electron tomography images showing nuclear envelope expansion in PfSUN2 inducible knock-down line comrade to NF54 and PfSUN1 inducible knock-down. N: Nucleus. Scale bar: 100 nm. **B.** Comparison of the maximum distance between the INM and ONM measured for the 3 lines. **C.** Comparison of the circularity index of trophozoite of the 3 lines. **D.** Comparison of the distribution of nuclear area (µm²) and **E.** Nuclear area to cell area ratio of the 3 lines. **F.** Comparison of the distribution of Food Vacuole (FV) area (µm²) and **G.** FV area to cell area ratio of the 3 lines. All measurements and morphometric analysis were performed using the measure feature in ImageJ. Error bars represent Standard Deviation. Statistical significances between different groups were determined using an unpaired student’s t-test (indicates *** P<0.001, ** P<0.01, * P<0.05, and ns; P>0.05).

**Figure 7.**
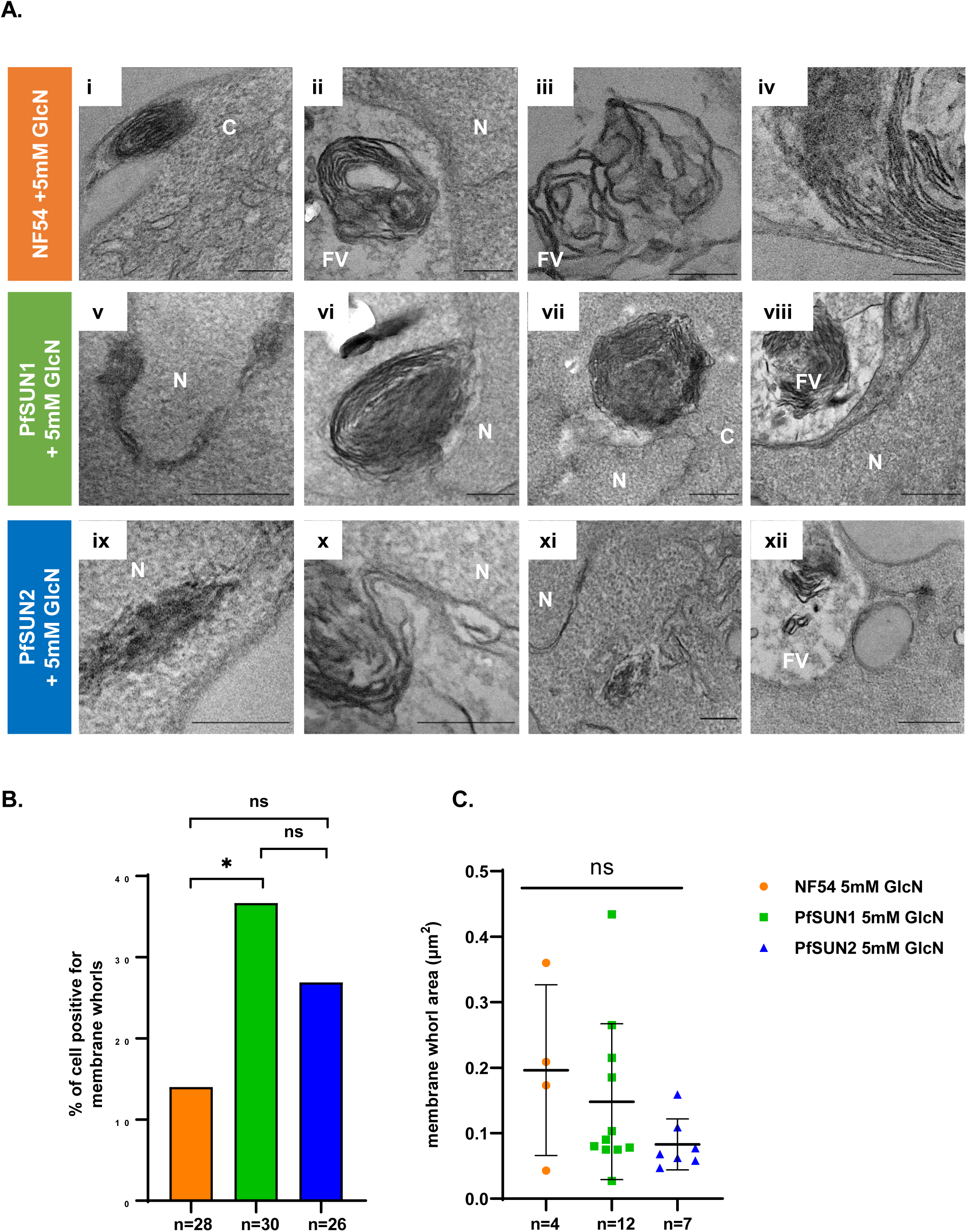
Loss of PfSUN1 and PfSUN2 expression results in nuclear membrane expansion, and membrane whorls formation. **A.** Ultrastructure analysis of late trophozoite stage of PfSUN1-HA-*glmS* and PfSUN2-HA-*glmS* grown on 5 mM GlcN for 82h was performed using TEM. NF54 wild type parasites grown on 5 mM GlcN for 82h were used as control (i-iv). Representative TEM images showing the expansion of the nuclear membrane and subsequent formation of membranous whorls (MW), that are later observed in the Food Vacuole (FV), following inducible knock-down of PfSUN1 (v-viii) and PfSUN2 (ix-xii)**. B.** Membrane whorls abundance and **C.** area distribution (µm²) in each of the 3 lines. All measurements were performed using the measure feature in ImageJ. Error bars represent Standard Deviation. Statistical significances between different groups were determined using an unpaired student’s t-test (indicates *** P<0.001, ** P<0.01, * P<0.05, and ns; P>0.05). Scale Bars: i, ii, iii, v, vi, vii, ix, x, xi 200nm; iv, viii and xii 500nm.

### PfSUN1 is essential for activating the DNA damage response

Previous work in mammalian cells showed that the mouse SUN domain proteins SUN1 and SUN2 play an important role in the DNA damage response (DDR) (29). Malaria parasites replicate their haploid genome multiple times through consecutive mitotic cycles and are particularly prone to error during DNA replication. Therefore, we were interested to determine whether PfSUN1 and PfSUN2 play a role in the *P. falciparum* DDR. To this end, we applied a DDR assay which was recently optimized for *Plasmodium spp.,* and uses anti γ-H2A.X antibody that recognizes the phosphorylated PfH2A, which serves as a marker for DNA damage in *P. falciparum* (30). The two transgenic lines, PfSUN1-HA-*glmS* and PfSUN2-HA-*glmS* were exposed to a sub lethal dose of X-ray irradiation (3000 rad) and the kinetics PfH2A phosphorylation was followed in presence or absence of PfSUN1 expression (**Fig. 8A**). We found that the PfSUN1-HA-*glmS* parasites, which were grown without GlcN, showed a decrease in the phosphorylation level of PfH2A over time (4 hours after irradiation), corresponding with the repair of DNA damage caused by irradiation (**Fig. 8A**, left panels). However, parasites grown on GlcN for 72h prior to the X-ray irradiation were unable to repair the DNA damage and the levels of PfH2A phosphorylation remained constant over time (**Fig. 8A**, right panels). Nonetheless, we show that GlcN had no effect on the ability of NF54 parasites to activate DDR (**Fig. 8B**). We were unable to detect any difference in DNA damage repair in the presence or absence of PfSUN2 (data not shown).

**Figure 8.**
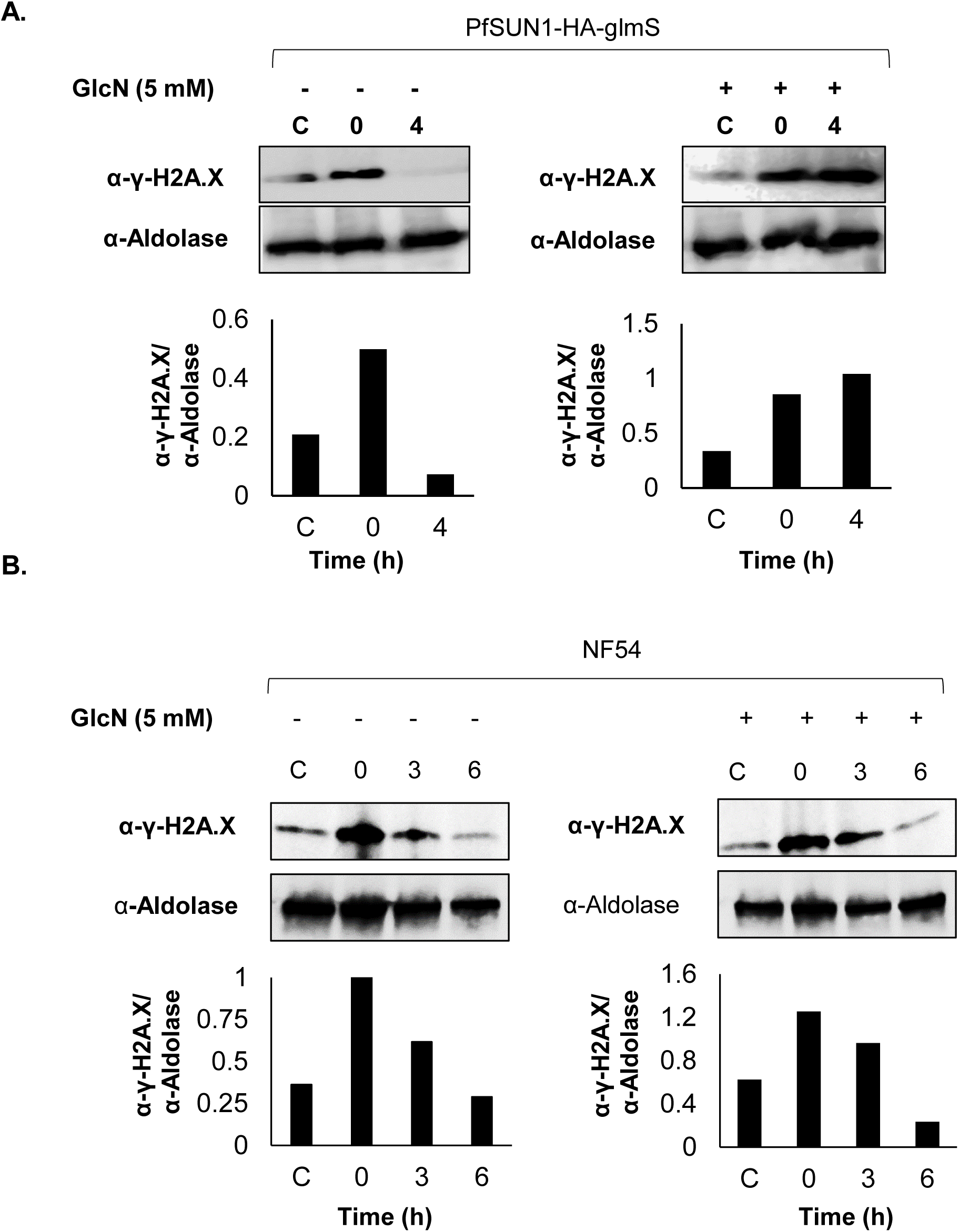
PfSUN1 is essential for activating the DNA damage response. **A.** The role of PfSUN1 and PfSUN2 in DNA damage repair by irradiating PfSUN1-HA-*glmS* and PfSUN2-HA-*glmS* parasites which were grown for 72h either on regular media (left) or on media supplemented with 5 mM GlcN (right). Tightly synchronized ring stages were exposed to a sub lethal dose of X-ray irradiation (6000 rad) **Upper panel:** Dynamic in phosphorylation of PfH2A measured by WB analysis of protein extracted from PfSUN1-HA-*glmS* parasites before and after irradiation in the presence (left) or absence (right) of PfSUN1 expression. C; Control untreated, 0; 15 minutes after X-ray irradiation and 4; 4 hours after irradiation. Activation of DDR and subsequent repair was analyzed using anti γ-PfH2A.X antibody. Anti-aldolase antibody was used for loading control. **Lower Panel:** Densitometry quantification of the ratio between the WB signals obtained for γ-PfH2A.X and aldolase presented above. Densitometry analysis was performed using ImageJ software. **B.** Similar experiment was performed on NF54 parasites grown for 72h either on regular media (left) or on media supplemented with 5 mM GlcN (right). Tightly synchronized ring stages were exposed to a sub lethal dose of X-ray irradiation (6000 rad). **Upper panel:** Dynamic in phosphorylation of PfH2A measured by WB analysis of protein extracted from control (untreated) and NF54 parasites before and after irradiation in the presence (right) or absence (left) of GlcN. C; Control untreated, 0; 15 minutes after X-ray irradiation, 3; 3 hours after irradiation and 6; 6hours after irradiation. Activation of DDR and subsequent repair was analyzed using anti γ-PfH2A.X antibody. Anti-aldolase antibody was used for loading control. **Lower Panel:** Densitometry quantification of the ratio between the WB signals obtained for γ-PfH2A.X and aldolase presented above. Densitometry analysis was performed using ImageJ software.

## Discussion

The complex biology of *Plasmodium* parasites and particularly their unique replication by closed mitosis during schizogony, requires special adaptations of their nuclear structure and functions that enable them to proliferate and thrive in different cellular niches within their hosts. Similar to higher eukaryotes, specific nuclear functions in *Plasmodium* were found to be linked to different sub-nuclear compartments. A marked example is their nuclear periphery, a sub-nuclear compartment that was implicated in regulation of gene expression, particularly of genes involved in virulence and immune evasion (3). Nonetheless, since only few components of the nuclear envelope have been identified, the nuclear periphery of *Plasmodium* refers primarily to the margins of the DNA staining imaged by light microscopy, while the precise morphological determination of this sub-nuclear compartment remains elusive. In addition, the role of the LINC complex that links the nuclear periphery with the cytoplasm remained understudied in these parasites, despite its importance for diverse nuclear functions, including nuclear replication and positioning, movement of chromosomes and DNA repair. Moreover, in model eukaryotes, some of the nuclear dynamics required for replication and chromosome movement were shown to involve interactions between the LINC complex and lamins (28) that serve as the nuclear scaffold below the inner nuclear membrane. Thus, in an organism such as *Plasmodium* that lacks conventional lamins, the mechanisms by which the LINC complex could mediate such nuclear dynamics remains a mystery, similar to African trypanosomes that do not encode for conventional lamins or canonical components of the LINC complex (31). SUN-domain proteins are major components of the LINC complex with many organisms expressing different SUN-domain variants in a cell cycle and in a tissue-specific manner (32). One example is the SUN-domain protein (TgSLP1) of the apicomplexan parasite *Toxoplasma gondii* that showed stage specific expression and was found to be essential for nuclear division and centrocone integrity (33). Interestingly, the two SUN-domain proteins which we found to be essential for the proper proliferation of blood stage *P. falciparum* parasites, have different temporal transcription profiles during the intra-erythrocytic development cycle (IDC). It appears that *pfsun1* transcripts are highly abundant in male gametocytes and ookinetes and transcript levels of *pfsun2* peak in mature liver and blood stage merozoites (34). These differential patterns may imply that in addition to their roles in the IDC they could also be important for parasite-specific functions in other stages of the parasite life cycle. Recent systematic screen for *Plasmodium* fertility genes identified *P. berghei* SUN1 complex, and demonstrated its essentiality for male gamete fertility by linking the microtubule organizing center to the nuclear envelope and enabling mitotic spindle formation during male gametogenesis (21).

While both PfSUN1 and PfSUN2 are important for parasite proliferation during the IDC, each could have specific functions as it appears that PfSUN2 is more tightly associated with heterochromatin while PfSUN1 plays a specific role in the DNA damage response. We have previously demonstrated that the *Plasmodium* NE is actively remodeled during asexual replication (25), changing the number, organization and the orientation of the nuclear pore complexes (NPC). Interestingly, the NPC clusters change their directionality as mitosis progresses indicating the possible rotation of the nucleus, which in some cases reaches up to 180 degrees during late schizogony (25). Remarkably, during this rotation the cluster of the NPC maintains its orientation facing the parasite’s membrane, providing evidence for possible cytoplasmic influence on NPC localization during the transition from a multinucleated syncytium to multiple autonomous cells. Interestingly, we observed similar directionality in the localization of both PfSUN1 and PfSUN2 throughout the IDC. PfSUN1 displays a highly polarized localization pattern in mature merozoites just before egress that is reminiscent of the previously observed clustering of NPCs at this stage. However, even though the NPC cluster at that stage is found adjacent to the foci of PfSUN1, they do not completely co-localize at any of the asexual stages, similar to what was observed in the fission yeast *S. cerevisiae,* where the SUN domain protein MPS3 does not co-localize with the NPC (35). Nonetheless, it was shown that the disruption of the LINC complex, as a critical component of force transmission between the nucleus and cytoskeleton, caused impaired nuclear positioning and cell polarization (36, 37). The dynamic nuclear positioning and the polarity of NPC clusters during plasmodium schizogony appear to be associated with the polarity of PfSUN1/2 accumulation observed here. Their cell-cycle-dependent polarity together with the slow growth rate and abnormal schizont morphology of parasites in which PfSUN1/2 were knocked-down, strongly suggests that PfSUN proteins play an important role in nuclear division and cellular segregation during schizogony. In the rodent malaria parasite *P. berghei*, PbSUN1 displayed a polarized localization near spindle poles, where it facilitates the rapid genome replication and NE remodeling in male gametocytes (22). These findings align with the polarity of PfSUN1 in segregating nuclei during asexual schizogony in *P. falciparum*, highlighting a possible conserved role across apicomplexan in coordinating nuclear-cytoskeletal interactions. Interestingly, *Cryptosporidium spp*. encode for orthologs of LINC complex components, which could also be implicated in its complex division processes that includes synchronized nuclear replication and membrane remodeling during sexual and asexual stages, however, this hypothesis requires further experimental evidence (38)

In addition to their impaired ability to segregate properly, and the observed morphological alteration of the perinuclear space, the most striking morphological phenotype was the appearance of the membrane whorls, which was more significant following PfSUN1 knocked-down. These membrane whorls appeared to be forming and pinching out at a distinct location at the NE and in many cases were finally observed at the parasite food vacuole - presumably for lysosomal-like degradation (39, 40). This could represent a sequential process that is reminiscent of microautophagy described in yeast, where ER whorls are translocated to the lysosome for degradation in mechanisms that are similar to known autophagy machinery (41). SUN-domain proteins were previously implicated in homeostasis of the nuclear envelope in *S. cerevisiae*, where mutations in the SUN protein MPS3 led to over proliferation of the NE (42), similar to what we observed here. It is interesting to note that depletion of SUN1 in *P. berghei* led to significant changes in lipid metabolism, linking NE dynamics with broader cellular processes such as membrane biogenesis and metabolic regulation (22). In addition, SUN proteins were shown to play a role in remodeling the NE, during open mitosis, by facilitating membrane removal from chromatin through NE breakdown (43). Evidently, disassembly of the LINC complex is required to control nuclear envelope dynamics during ER stress, to allow specific autophagy responses that maintain the homeostasis of the NE and the ER (44). Altogether these data suggest that SUN proteins in *P. falciparum* may play a role in the homeostasis of the NE and the ER.

In addition to the abnormal replication phenotype, we found that PfSUN1 is essential for activating the DNA damage response (DDR). In recent years there is mounting evidence that the LINC complex plays an important role in different aspects of the DDR (15). Recently we showed that a plasmodium *bona fide* SR protein, PfSR1, is essential for DNA repair that takes place at specific foci at the nuclear periphery (45). We found that after exposing the parasite to a source of damage, PfSR1 is recruited to the site of damage where it co-localizes with PfRAD51. Furthermore, we demonstrated that PfSR1 expression is essential for the recruitment of PfRAD51 to the site of damage as well as for the parasite’s ability to activate DDR. These data suggest that the nuclear periphery of *P. falciparum* contains functional sub-compartments specific for DNA repair. In *C. elegans* as well as in mammalian cells, SUN-domain proteins were shown to be involved in recruitment of DNA repair proteins such as DNA-dependent protein kinase (DNA-PK), Rad51, Ku70/Ku80 and FAN1 nuclease to the site of damage. In addition, SUN domain proteins can influence which repair mechanisms will be applied by promoting HR over NHEJ (14). Damaged DNA must be mobile in the nucleoplasm to be directed and reach the site of repair at the nuclear periphery. It was shown that chromatin surrounding a double strand break has greater mobility which is mediated by cytoplasmic microtubules that interact with nucleoplasmic 53BP1 through required interaction with SUN domain proteins (13). Although the exact mechanistic role of components of the LINC complex in DDR remains elusive, it is reasonable to suggest that SUN-domain proteins in plasmodium could contribute to the formation of a functional repair site at the nuclear periphery and mobilize both the damaged locus and components of the repair machinery to create the sub-nuclear repair site.

Our study provides the first glimpse into the role of SUN proteins as possible components of the *Plasmodium* LINC complex essential for various nuclear functions in blood stage parasites. Identification of PfSUN proteins could further be exploited for the discovery of other components of the LINC complex and the nuclear envelope. In addition, this knowledge could be further utilized to investigate their involvement in the dynamics of gene positioning at the nuclear periphery, interactions and movement of chromosomes and anchoring specific loci to functional sub-nuclear sites.

## Materials and Methods

### Parasite culturing and parasitemia counts

Parasites were kept in continuous culture as in (46). All parasites used were derivatives of the NF54 parasite line and were cultivated at 5% haematocrit in RPMI 1640 medium, 0.5% Albumax II (Invitrogen), 0.25% sodium bicarbonate, and 0.1 mg/ml gentamicin. Parasites were incubated at 37°C in an atmosphere of 5% oxygen, 5% carbon dioxide, and 90% nitrogen. Depending on required experiments, parasitemia (% of iRBCs out of the total RBCs) was kept at 0.1 - 5 % by dilution, and the medium was changed at least every second day. Parasite cultures were synchronized using percoll/sorbitol gradient centrifugation as previously described (47, 48). Briefly, infected RBCs were layered on a step gradient of 40%/70% percoll containing 6% sorbitol. The gradients were then centrifuged at 12,000g for 20 minutes at room temperature. Highly synchronized, late-stage parasites were recovered from the 40%/70% interphase, washed twice with complete culture media, and placed back in culture with an adjusted haematocrit of 5% Parasite cultures were synchronized using sorbitol as previously described (49), briefly parasite culture the culture was transferred into a 15 ml falcon tube and spun down at 1800 x g for 3 minutes. The supernatant was discarded and the pellet was resuspended in a pre-warmed 5 % D-sorbitol solution. After incubation for 10 minutes at 37 °C, the falcon tube vortexed for 5 seconds and centrifuged by 800 g for 5 minutes. The pellet was washed with RPMI complete medium and re-cultured in fresh RPMI complete medium. The haematocrit was adjusted to 5 %. Sorbitol synchronization leads to a culture holding only ring stage parasites. Transgenic parasites were cultured in the presence of drugs corresponding to the respective selectable marker used. The level of parasitemia was calculated either by Flow Cytometry. For flow cytometry, aliquots of 50µl parasite cultures were washed in PBS and incubated for 30 min with 1:10,000 SYBR Green I DNA stain (Life Technologies). The fluorescence profiles of infected erythrocytes were measured on CytoFLEX (Beckman Coulter) and analysed by the CytExpert software.

### Parasite transfection

Parasites were transfected as described (50). Briefly, 0.2 cm electroporation cuvettes were loaded with 0.175ml of erythrocytes and 50-100 µg of purified plasmid DNA in incomplete cytomix (8.95g KCl, 0.017g CaCl, 0.76g EGTA,1.02 g MgC2, 0.871g K2HPO4, 0.68g KH2PO4, 7.08 g HEPES) solution. Electroporation was performed using the Gene Pulser Xcell (Bio-Rad, 310 V, 950 µF, ∞ Ω). Electrophorized blood was washed twice in complete RPMI medium, transferred to a fresh culture flask with 10ml of fresh complete RPMI, schizonts isolated using percoll/sorbitol synchronization and blood to an adjusted haematocrit of 5%. After 24 hours the selection drug was added and medium was changed daily for the first 5 days and every second day from that day onwards. Stable transfectants carrying plasmids with an hDHFR-selectable marker were selected on 4 nM WR99210 and those carrying yDHODH were selected on 1.5 µM DSM1. Stable transfectants carrying plasmids with BSD-selectable marker were initially selected on 2 μg/ml blasticidin-S (Invitrogen). In order to obtain parasites carrying large plasmid copy numbers, these cultures were then subjected to elevated concentrations of 6–10 µg/ml blasticidin-S, depending on experimental design.

### Bioinformatics analyses

In order to find putative SUN domain proteins in *P. falciparum* a BLAST search was performed against the SUN domain annotated sequence of Human SUN domains of Human sun domain proteins in the Plasmodium database PlasmoDB (51).Illustrations of domain architecture were generated using IBS illustrator (52). The SUN domain sequences of selected SUN domain proteins were obtained from UniProt (53). Membrane topology prediction was analyzed using Protter (54) Multiple-sequence alignment of PfSUN1 and PfSUN2 (PF3D7_1215100 and PF3D7_1439300) was performed using CLUSTALO (55) and further analyzed using ESPript3 program (56). A homology model of the SUN domain of PfSUN1 and PfSUN2 was built by comparative modeling using the crystal structure of HsSUN2 (Protein Data Bank entry 4DXT,) by using the I-TASSER server (57). The structure visualization of the PfSUN1 and PfSUN2 three-dimensional (3D) model was performed using the PyMOL program (58). Gene expression profiles were analyzed using resources available in the PlasmoDB database, derived from previously published RNA-seq datasets. The datasets were pre-normalized to Transcripts Per Million (TPM) by the original authors, allowing for direct comparison. For the IDC, only well annotated data points were included to ensure accuracy. *P. berghei* data were used for liver stages due to the lack of P. falciparum data, enabling a complete life cycle profile. Mean TPM values were averaged from the combined datasets to provide a reliable representation of gene expression (59–70).

### Immunofluorescence assay

IFA was performed as described (30). In brief, iRBCs were washed twice with PBS and resuspended in freshly prepared fixative solution, 4% paraformaldehyde [EMS; Electron Microscopy Sciences] and 0.0075% glutaraldehyde [EMS] in PBS, for 30 min at room temperature. Following fixation, iRBCs were permeabilized with 0.1% Triton X-100 (Sigma) in PBS and then blocked with 3% bovine serum albumin (BSA; Sigma) in PBS. Alternatively, parasites were released from iRBCs with saponin, washed and fixed in 4% PFA on ice for 1 hour. Cells were then incubated with the following primary antibodies, used at the indicated dilutions: mouse anti-GFP (Roche 1:300), mouse anti-HA (Roche:100), rabbit anti-myc (Cell Signalling 1:100), for ER staining rabbit anti BiP (MR4 1246, 1:200) for 1.5 h at room temperature and washed three times in PBS. Following this, cells were incubated with the following secondary antibodies conjugated to Alexa fluorophores: Alexa Fluor 488 goat anti-mouse, Alexa Fluor 568 goat anti-mouse, Alexa Fluor 568 goat anti-rabbit (Life Technologies; 1:500) antibodies for 1 h at room temperature. Cells were washed three times in PBS and laid on polytetrafluoroethylene (PTFE) printed slides (EMS) and mounted in ProLong Gold antifade reagent with DAPI (Molecular Probes). Fluorescent images were obtained using a Plan Apo λ 100× oil immersion lens (numerical aperture [NA] = 1.5; working distance [WD] = 130 μm) on a Nikon Eclipse Ti-E microscope equipped with a CoolSNAPMyo CCD camera. Images were processed using NIS-Elements AR (4.40 version) software.

### Immunogold labeling and electron microscopy analysis

Cells were fixed in 4% paraformaldehyde with 0.1% glutaraldehyde in 0.1M cacodylate buffer (pH = 7.4) for 1 hour at room temperature and kept overnight at 4°. The samples were soaked overnight in 2.3M sucrose and rapidly frozen in liquid nitrogen. Frozen ultrathin (70–90 nm) sections were cut with a diamond knife at −120°C on a Leica EM UC6 ultramicrotome. The sections were collected on 200-mesh Formvar coated nickel grids. Sections were blocked with a solution containing 1% BSA, 0.1% glycine, 0.1% gelatin, and 1% Tween 20. Immuno-labeling was performed using affinity purified anti-GFP antibodies (1:20, Abcam), overnight at 4°C, followed by exposure to goat anti-Rabbit IgG coupled to 10-nm gold particles (1∶20, Jackson ImmunoResearch), for 30 min at room temperature. Contrast staining and embedding were performed as previously described (71). The embedded sections were viewed and photographed with a FEI Tecnai SPIRIT (FEI, Eidhoven, Netherlands) transmission electron microscope operated at 120 kV, and equipped with an EAGLE CCD Camera.

### Transmission electron microscopy

iRBCs were pelleted, washed twice with PBS and fixed using 2% formaldehyde and 2.5% glutaraldehyde in 0.1 M Cacodylate buffer, pH 7.4 at room temperature for 2h, following an overnight incubation at 4°C. The iRBCs were washed and post-fixed with 1% osmium tetroxide in the same buffer for 1 h at room temperature, the samples were dehydrated in graded ethanol series and embedded in Epon. Thin sections (70–90 nm) were prepared using an Ultracut UCT microtome (Leica). Followed post staining with 2% uranylacetate and Reynold’s lead citrate, and viewed in a Jeol^®^ JEM-1400 Plus TEM (Jeol^®^, Tokyo, Japan) operating at 100 kV, equipped with ORIUS SC600 CCD camera (Gatan^®^, Abingdon, United Kingdom), and Gatan Microscopy Suite program (DigitalMicrograph, Gatan^®^, UK). Image processing was performed on the DM3. files using FIJI (72)

### STORM imaging and analysis

Stochastic Optical Reconstruction Microscopy (STORM) imaging was performed as described (73) in brief Parasite cultures were saponin lysed and washed three times with fresh PBS. Parasites were then re-suspended on ice with fresh fixative solution, 4% Paraformaldehyde (EMS) in PBS for 1h. Fixed parasites were allowed to air dry in an 8 well-chambered cover glass system 1.5H (In Vitro Scientific) for 2-3 hours at room temperature and covered with glycine (125 mM glycine in PBS) to quench unreacted aldehyde groups. Samples were washed three times with PBS and permeabilized with 0.1% Triton-x 100 (PBS) for 10 min at room temperature, washed with PBS and blocked with 3% BSA (IgG free, Sigma) in 0.025% Tx-100/PBS for 1 hour at room temperature. Primary antibodies, mouse anti-GFP (1:300, Roche), mouse anti-HA (1:100 Roche), rabbit anti-Halo (1:300, Promega), rabbit anti-H3K9me3 (1:300 Abcam) and rabbit anti-H3K9Ac (1:300 Abcam) were diluted in 3% BSA containing 0.025% Tx-100 in PBS and applied for 1 hour at room temperature, followed by 5 washes in PBS containing 0.025% Tx-100. Secondary antibodies, Goat anti-mouse Alexa647 and Goat anti-Rabbit Alexa568 (1:500, life technologies) were applied for 1 hour in 3% BSA containing 0.025% in PBS at room temperature, followed by 5 washes with 0.025% Tx-100 in PBS. Parasite nuclei were labeled with YOYO-1 (1:1000, life technologies) for 20 min at room temperature, followed by 3 washes with PBS. STORM was performed by a Nikon Eclipse Ti-E microscope with a CFI Apo TIRF × 100 DIC N2 oil objective (NA 1.49, WD 0.12 mm). Multi-channel calibration was performed prior data acquisition using fluorescent TetraSpeckTM microspheres 0.1μm in diameter (Life-technologies, Molecular Probes). Stained cells were placed in Glox-MEA imaging buffer containing 50mM MEA, 10% glucose (D2O), Glox (11.2mg/ml Glucose Oxidase (Sigma) and 1.8mg/ml Catalase (Sigma, C30-500MG)) in dilution buffer (50mM NaCl, 200mM Tris in D2O). Samples were illuminated by 561nm and 647nm excitation lasers in changing intensity over the duration of the imaging sequence (typically, using 50-100% power). 488nm laser was used at 0.5% power to visualizenuclei by TIRF. For each acquisition, 10000 frames were recorded onto a 256×256-pixel region (pixel size 160nm) of an Andor iXon-897 EMCCD camera. Super-resolution images were reconstructed from a series of the least 5000 images per channel using the N-STORM analysis module, version 1.1.21 of NIS Elements AR v. 4.40 (Laboratory imaging s.r.o.).

### Plasmid construction

In order to express PfSUN1 fused to a GFP tag at its’ C’ terminus, the genomic sequence of PF3D7_1215100 was amplified using PfSUN1-F 5’-AACTGCAGAAAAATGAACATAAGCAACAGT-3’ and PfSUN1-R 5’-GAAGATCTAAATTTTCTTATACATCTTTTT-3’ and cloned into the pHTIDH expression vector (24) using PstI and BglII to generate pHTIDH-SUN1GFP. In order to express PfSec13 fused to a Halo tag, the genomic sequence of Pf3D7_1230700 was amplified from using the primers Sec13Halo-F GCGATCGCCATGAACGAATTAGTAGTG and Sec13Halo-R CTGCAGATATTGTTCATATGTGTATT and cloned into the pHBIcHalo (45) expression vector using PstI and AsisI to generate pHBIPfSec13cHalo.

The plasmid pVSUN1-myc was generated in two steps: The plasmid pVB-myc (73) was used as template, first the selection marker was changed to BSD using the primers BSD-F (5’-GCGGCCGCAAAATGCCTTTGTCTCAAG-3’) and Hrp23’-R (5’-AACCAACGCGTTGGTGCAGTTTAAT-3’), the sequence was amplified from pVBh (74) and inserted between the NotI and BstXI sites to generate the pV-myc construct. As a second step, PfSUN1 genomic sequence was amplified using SUN1HpaI-F (5’-AGTTAACATGAACATAAGCAACAG-3’) and SUN1KasI-R (5’-TTGGCGCCTTTAAATTTTCTTATAC-3’). The SUN1 1-190 fragment was amplified using SUN1HpaI-F and SUN1-190KasI-R (5’-GGCGCCTATTATTATTATTATTATATAGTGTAATAAATCCTG-3’). The SUN1Δ250-600 sequence was ordered as a gene block in pUC-57 (IDT) and cloned into pV-myc construct with HpaI and KasI. PCR amplifications were performed using high-fidelity PrimeSTAR HS DNA polymerase (Takara Bio) or Q5 high fidelity polymerase (New England Biolabs; NEB) utilizing NF54 genomic DNA as template, unless otherwise specified.

### Generation of transgenic lines and inducible knock-down

The PfSUN1-HA-glms and PfSUN2-HA-glms lines were created using the plasmid pSLI-HAx3glmS (75) as backbone. pSLI-HAx3glms was generated on the basis of the pSLI-2 X FKBP-GFP construct (27). The homology region for PfSUN1 (PF3D7_1215100) was cloned using the primers SUN1HR-Long-F (5’-GCGGCCGCTAATGGGCACTTGAATCT-3’) and SUN1-3UTR-R (5’-CTATATATTTTTGTTGGTACCACATG-3’) (NotI and XmaI). The homology region of PfSUN2 (PF3D7_1439300) was cloned using the primers SUN2HR-Long-F (5’-GCGGCCGCTAAACGTTAGTTGATAAAATAAAAACTATCG-3’) and SUN2-3UTR-R2 (5’-TAATAACTGTTGTGAAAAGTTTCTAACAGC-3’) (NotI and XmaI).

Analysis of the integrated construct was performed using diagnostic PCR at the integration sites followed by sequencing and southern blots of selected subclones.

For inducible knockdown, PfSUN1-HA-glmS and PfSUN2-HA-glmS parasites were grown in the presence or absence of 5 mM D-(+)-Glucosamine hydrochloride (GlcN) (Sigma) over a 72h time course. Following treatment parasites were released by saponin lysis and sequentially analyzed in western blot.

### Western blotting

To collect parasite proteins, infected RBCs were lysed with saponin, and parasites were pelleted down by centrifugation. The parasite pellet was subsequently washed twice with PBS and lysed in 2× Laemmli sample buffer or Urea/SDS lysis buffer. The protein lysates were centrifuged and the supernatants were subjected to SDS-PAGE (gradient, 4 to 20%; Bio-Rad) and electroblotted onto a nitrocellulose membrane. Immunodetection was carried out by using mouse anti-GFP (1:1000, Roche), mouse anti-HA primary antibody (1:1000, Roche), rabbit anti-γH2A.X antibody (1:1000, Cell signaling) and rabbit anti-aldolase (1:3000, Abcam). The secondary antibodies used were antibodies conjugated to horseradish peroxidase (HRP), goat anti-rabbit (1:10000, Jackson ImmunoResearch). The immunoblots were developed in EZ/ECL solution (Israel Biological Industries).

### X-ray irradiation of parasites and DNA repair assay

DNA damage of parasites and repair assay was performed as previously described (30). Briefly, tightly synchronized ring stage parasites were exposed to 6000 rads X-ray irradiation using a PXi precision X-ray irradiator set at 225 kV and 13.28 mA. Immediately following irradiation, parasites were put back to culture to allow them to repair the damaged DNA. Protein was extracted 15 min after irradiation (0 h), as well as from parasites collected at 4 h after irradiation. Proteins extracted from untreated iRBCs were used as control. Western blot analysis was used to follow the changes in γ-*Pf*H2A compared with the housekeeping control gene aldolase in each treatment. These western blots were subjected to densitometry analysis to calculate the ratio between γ-*Pf*H2A levels and aldolase.

### Southern Blot

Southern blots were performed as described in (24). Briefly, genomic DNA isolated from recombinant parasites was digested to completion by the restriction enzymes XbaI and BglII (PfSUN1-HA-glms) and XbaI and SacI (PfSUN2-HA-glms) (NEB) and subjected to gel electrophoresis using 1% Agarose in Tris/Borate/EDTA (TBE). The DNA was transferred to high-bond nitrocellulose membrane by capillary action after alkaline denaturation. DNA detection was performed using DIG High Prime DNA Labeling and Detection starter kit (Roche). The HR sequences were amplified from pSLI-SUN1-HAx3-glms and pSLI-SUN2-HAx3-glms, DIG labelled and used as probes.

## Acknowledgments

This work was supported partially by the Israeli Academy for Science, Israel Science Foundation (ISF) Grant 409/23 and in part by the United States − Israel Binational Science Foundation (BSF) grant 2023133 and European Research Council (erc.europa.eu) Consolidator Grant 615412 (to R.D.). RD is also supported by Ministry of Science and Technology Grant 103240; the United States − Israel Binational Science Foundation (BSF) grant 2019236, and the Dr. Louis M. Leland and Ruth M. Leland Chair in Infectious Diseases.

## References

WHO (2021) World Malaria Report, 2021.263.

Bozdech Z, et al. (2003) The transcriptome of the intraerythrocytic developmental cycle of Plasmodium falciparum. PLoS Biol 1(1):E5.

Deitsch KW & Dzikowski R (2017) Variant Gene Expression and Antigenic Variation by Malaria Parasites. Annu Rev Microbiol.

D’Angelo MA & Hetzer MW (2006) The role of the nuclear envelope in cellular organization. Cell Mol Life Sci 63(3):316–332.

Hetzer MW (2010) The nuclear envelope. Cold Spring Harb Perspect Biol 2(3):a000539.

Ungricht R & Kutay U (2017) Mechanisms and functions of nuclear envelope remodelling. Nat Rev Mol Cell Biol 18(4):229–245.

Zuleger N, Robson MI, & Schirmer EC (2011) The nuclear envelope as a chromatin organizer. Nucleus 2(5):339–349.

Bouzid T, et al. (2019) The LINC complex, mechanotransduction, and mesenchymal stem cell function and fate. J Biol Eng 13:68.

Mejat A & Misteli T (2010) LINC complexes in health and disease. Nucleus 1(1):40–52.

Razafsky D & Hodzic D (2009) Bringing KASH under the SUN: the many faces of nucleo-cytoskeletal connections. J Cell Biol 186(4):461–472.

Rothballer A, Schwartz TU, & Kutay U (2013) LINCing complex functions at the nuclear envelope: what the molecular architecture of the LINC complex can reveal about its function. Nucleus 4(1):29–36.

Tzur YB, Wilson KL, & Gruenbaum Y (2006) SUN-domain proteins: ’Velcro’ that links the nucleoskeleton to the cytoskeleton. Nat Rev Mol Cell Biol 7(10):782–788.

Lottersberger F, Karssemeijer RA, Dimitrova N, & de Lange T (2015) 53BP1 and the LINC Complex Promote Microtubule-Dependent DSB Mobility and DNA Repair. Cell 163(4):880–893.

Lawrence KS, et al. (2016) LINC complexes promote homologous recombination in part through inhibition of nonhomologous end joining. J Cell Biol 215(6):801–821.

Lambert MW (2019) The functional importance of lamins, actin, myosin, spectrin and the LINC complex in DNA repair. Exp Biol Med (Maywood) 244(15):1382–1406.

Liu Q, et al. (2007) Functional association of Sun1 with nuclear pore complexes. J Cell Biol 178(5):785–798.

Bupp JM, Martin AE, Stensrud ES, & Jaspersen SL (2007) Telomere anchoring at the nuclear periphery requires the budding yeast Sad1-UNC-84 domain protein Mps3. J Cell Biol 179(5):845–854.

Starr DA (2009) A nuclear-envelope bridge positions nuclei and moves chromosomes. J Cell Sci 122(Pt 5):577–586.

Schober H, Ferreira H, Kalck V, Gehlen LR, & Gasser SM (2009) Yeast telomerase and the SUN domain protein Mps3 anchor telomeres and repress subtelomeric recombination. Genes Dev 23(8):928–938.

Piccus R & Brayson D (2020) The nuclear envelope: LINCing tissue mechanics to genome regulation in cardiac and skeletal muscle. Biol Lett 16(7):20200302.

Sayers C, et al. (2024) Systematic screens for fertility genes essential for malaria parasite transmission reveal conserved aspects of sex in a divergent eukaryote. Cell Syst 15(11):1075–1091 e1076.

Zeeshan M, et al. (2024) A novel SUN1-ALLAN complex coordinates segregation of the bipartite MTOC across the nuclear envelope during rapid closed mitosis in Plasmodium. bioRxiv:2024.2012.2004.625416.

Aurrecoechea C, et al. (2009) PlasmoDB: a functional genomic database for malaria parasites. Nucleic Acids Res 37(Database issue):D539-543.

Dahan-Pasternak N, et al. (2013) PfSec13 is an unusual chromatin-associated nucleoporin of Plasmodium falciparum that is essential for parasite proliferation in human erythrocytes. Journal of cell science 126(14):3055–3069.

Weiner A, et al. (2011) 3D nuclear architecture reveals coupled cell cycle dynamics of chromatin and nuclear pores in the malaria parasite Plasmodium falciparum. Cell Microbiol.

Deitsch K, et al. (2007) Mechanisms of gene regulation in Plasmodium. Am J Trop Med Hyg 77(2):201–208.

Birnbaum J, et al. (2017) A genetic system to study Plasmodium falciparum protein function. Nat Methods 14(4):450–456.

Horn HF (2014) LINC complex proteins in development and disease. Curr Top Dev Biol 109:287–321.

Lei K, et al. (2012) Inner nuclear envelope proteins SUN1 and SUN2 play a prominent role in the DNA damage response. Curr Biol 22(17):1609–1615.

Goyal M, et al. (2021) Phosphorylation of the Canonical Histone H2A Marks Foci of Damaged DNA in Malaria Parasites. mSphere 6(1):e01131–01120.

Field MC (2023) Deviating from the norm: Nuclear organisation in trypanosomes. Curr Opin Cell Biol 85:102234.

Hieda M (2017) Implications for Diverse Functions of the LINC Complexes Based on the Structure. Cells 6(1).

Wagner M, Song Y, Jimenez-Ruiz E, Hartle S, & Meissner M (2023) The SUN-like protein TgSLP1 is essential for nuclear division in the apicomplexan parasite Toxoplasma gondii. J Cell Sci 136(21).

Amos B, et al. (2022) VEuPathDB: the eukaryotic pathogen, vector and host bioinformatics resource center. Nucleic Acids Res 50(D1):D898–D911.

Horigome C, Okada T, Shimazu K, Gasser SM, & Mizuta K (2011) Ribosome biogenesis factors bind a nuclear envelope SUN domain protein to cluster yeast telomeres. EMBO J 30(18):3799–3811.

Lombardi ML, et al. (2011) The interaction between nesprins and sun proteins at the nuclear envelope is critical for force transmission between the nucleus and cytoskeleton. J Biol Chem 286(30):26743–26753.

Lee YL & Burke B (2018) LINC complexes and nuclear positioning. Semin Cell Dev Biol 82:67–76.

Guerin A & Striepen B (2020) The Biology of the Intestinal Intracellular Parasite Cryptosporidium. Cell Host Microbe 28(4):509–515.

Matz JM (2022) Plasmodium’s bottomless pit: properties and functions of the malaria parasite’s digestive vacuole. Trends Parasitol 38(7):525–543.

Wiser MF (2024) The Digestive Vacuole of the Malaria Parasite: A Specialized Lysosome. Pathogens 13(3).

Schafer JA, et al. (2020) ESCRT machinery mediates selective microautophagy of endoplasmic reticulum in yeast. EMBO J 39(2):e102586.

Friederichs JM, et al. (2011) The SUN protein Mps3 is required for spindle pole body insertion into the nuclear membrane and nuclear envelope homeostasis. PLoS Genet 7(11):e1002365.

Turgay Y, et al. (2014) SUN proteins facilitate the removal of membranes from chromatin during nuclear envelope breakdown. J Cell Biol 204(7):1099–1109.

Kucinska MK & Molinari M (2023) Control of nuclear envelope dynamics during acute ER stress by LINC complexes disassembly and selective, asymmetric autophagy of the outer nuclear membrane. Autophagy:1–3.

Goyal M, et al. (2021) An SR protein is essential for activating DNA repair in malaria parasites. J Cell Sci 134(16).

Trager W & Jensen JB (1976) Human malaria parasites in continuous culture. Science 193:673–675.

Aley SB, Sherwood JA, & Howard RJ (1984) Knob-positive and knob-negative Plasmodium falciparum differ in expression of a strain-specific malarial antigen on the surface of infected erythrocytes. J Exp Med 160(5):1585–1590.

Calderwood MS, Gannoun-Zaki L, Wellems TE, & Deitsch KW (2003) Plasmodium falciparum var genes are regulated by two regions with separate promoters, one upstream of the coding region and a second within the intron. J Biol Chem 278(36):34125–34132.

Lambros C & Vanderberg JP (1979) Synchronization of Plasmodium falciparum erythrocytic stages in culture. J Parasitol 65(3):418–420.

Deitsch K, Driskill C, & Wellems T (2001) Transformation of malaria parasites by the spontaneous uptake and expression of DNA from human erythrocytes. Nucleic Acids Res 29(3):850–853.

Anonymous (2001) PlasmoDB: An integrative database of the Plasmodium falciparum genome. Tools for accessing and analyzing finished and unfinished sequence data. The Plasmodium Genome Database Collaborative. Nucleic Acids Res 29(1):66–69.

Liu W, et al. (2015) IBS: an illustrator for the presentation and visualization of biological sequences. Bioinformatics 31(20):3359–3361.

UniProt C (2015) UniProt: a hub for protein information. Nucleic Acids Res 43(Database issue):D204–212.

Omasits U, Ahrens CH, Müller S, & Wollscheid B (2014) Protter: interactive protein feature visualization and integration with experimental proteomic data. Bioinformatics 30(6):884–886.

Sievers F & Higgins DG (2014) Clustal Omega, accurate alignment of very large numbers of sequences. Multiple sequence alignment methods, (Springer), pp 105–116.

Gouet P, Robert X, & Courcelle E (2003) ESPript/ENDscript: Extracting and rendering sequence and 3D information from atomic structures of proteins. Nucleic Acids Res 31(13):3320–3323.

Yang J, et al. (2015) The I-TASSER Suite: protein structure and function prediction. Nat Methods 12(1):7–8.

Schrödinger L & DeLano W (2020) PyMOL. The PyMOL Molecular Graphics System, Version 2.

Toro-Moreno M, Sylvester K, Srivastava T, Posfai D, & Derbyshire ER (2020) RNA-Seq Analysis Illuminates the Early Stages of Plasmodium Liver Infection. mBio 11(1).

Chappell L, et al. (2020) Refining the transcriptome of the human malaria parasite Plasmodium falciparum using amplification-free RNA-seq. BMC Genomics 21(1):395.

Lindner SE, et al. (2019) Transcriptomics and proteomics reveal two waves of translational repression during the maturation of malaria parasite sporozoites. Nat Commun 10(1):4964.

Caldelari R, et al. (2019) Transcriptome analysis of Plasmodium berghei during exo-erythrocytic development. Malar J 18(1):330.

Toenhake CG, et al. (2018) Chromatin Accessibility-Based Characterization of the Gene Regulatory Network Underlying Plasmodium falciparum Blood-Stage Development. Cell Host Microbe 23(4):557–569 e559.

Gomez-Diaz E, et al. (2017) Epigenetic regulation of Plasmodium falciparum clonally variant gene expression during development in Anopheles gambiae. Scientific reports 7:40655.

Zanghi G, et al. (2018) A Specific PfEMP1 Is Expressed in P. falciparum Sporozoites and Plays a Role in Hepatocyte Infection. Cell reports 22(11):2951–2963.

Lopez-Barragan MJ, et al. (2011) Directional gene expression and antisense transcripts in sexual and asexual stages of Plasmodium falciparum. BMC Genomics 12:587.

Bartfai R, et al. (2010) H2A.Z demarcates intergenic regions of the plasmodium falciparum epigenome that are dynamically marked by H3K9ac and H3K4me3. PLoS Pathog 6(12):e1001223.

Wichers JS, et al. (2019) Dissecting the Gene Expression, Localization, Membrane Topology, and Function of the Plasmodium falciparum STEVOR Protein Family. mBio 10(4).

Otto TD, et al. (2010) New insights into the blood-stage transcriptome of Plasmodium falciparum using RNA-Seq. Mol Microbiol 76(1):12–24.

Lasonder E, et al. (2016) Integrated transcriptomic and proteomic analyses of P. falciparum gametocytes: molecular insight into sex-specific processes and translational repression. Nucleic Acids Res 44(13):6087–6101.

Tokuyasu KT (1986) Application of cryoultramicrotomy to immunocytochemistry. J Microsc 143(Pt 2):139–149.

Schindelin J, et al. (2012) Fiji: An open-source platform for biological-image analysis. Nature Methods 9:676–682.

Fastman Y, et al. (2018) An upstream open reading frame (uORF) signals for cellular localization of the virulence factor implicated in pregnancy associated malaria. Nucleic Acids Res 46(10):4919–4932.

Dzikowski R & Deitsch KW (2008) Active Transcription is Required for Maintenance of Epigenetic Memory in the Malaria Parasite Plasmodium falciparum. J Mol Biol.

Heinberg A, et al. (2022) A nuclear redox sensor modulates gene activation and var switching in Plasmodium falciparum. Proc Natl Acad Sci U S A 119(33):e2201247119.

