## Supplementary figures and legends for "SUN-domain proteins of the malaria parasite *Plasmodium falciparum* are essential for proper nuclear division and DNA repair"

**Figure S1.**

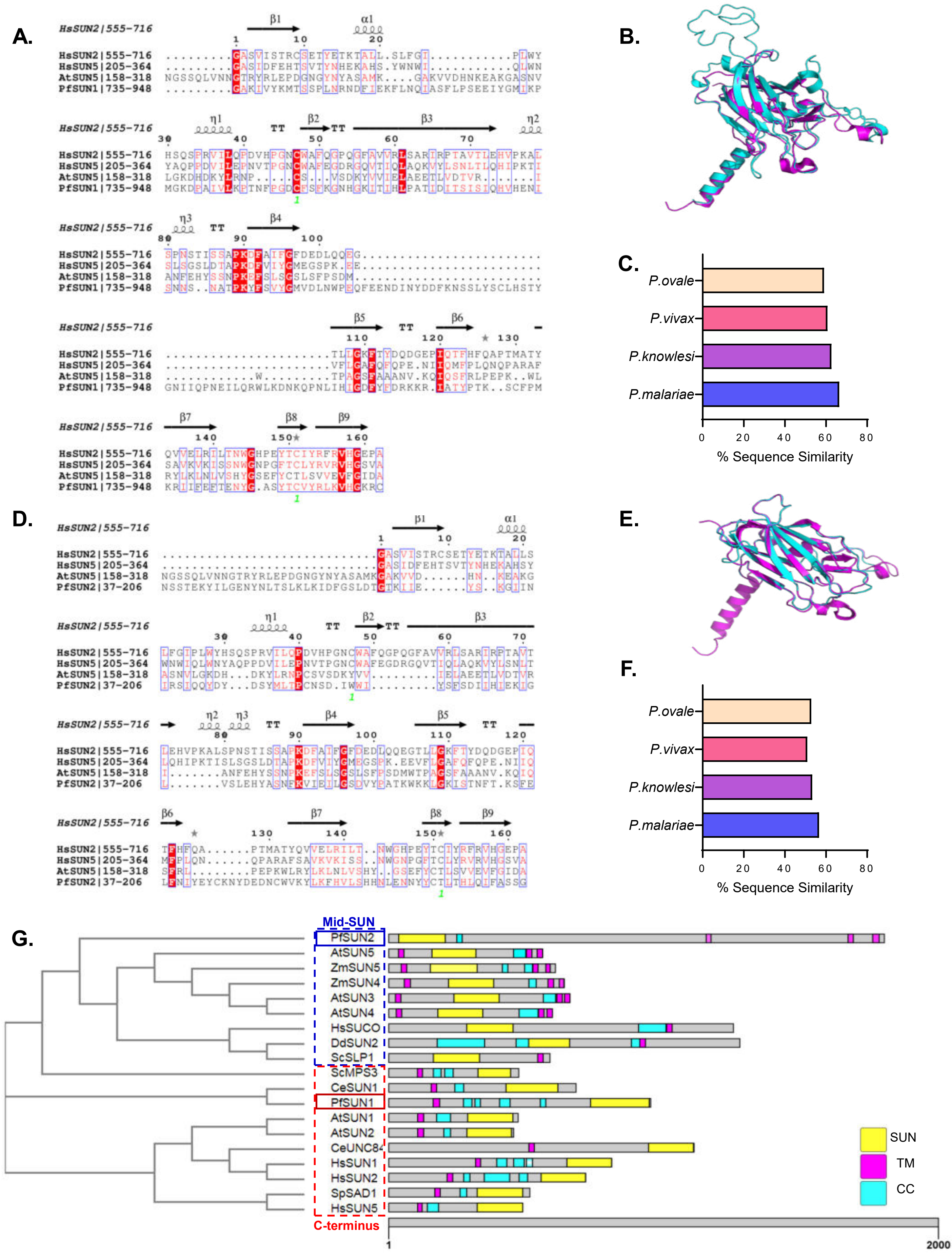

A.
HsSUN2|555-716
HsSUN5|205-364
AtSUN5|158-318
PfSUN1|735-948
B.
C.
P.ovale
P.vivax
P.knowlesi
P.malariae
% Sequence Similarity
D.
HsSUN2|555-716
HsSUN5|205-364
AtSUN5|158-318
PfSUN2|37-206
E.
F.
P.ovale
P.vivax
P.knowlesi
P.malariae
% Sequence Similarity
G.
Mid-SUN
PfSUN2
AtSUN5
ZmSUN5
ZmSUN4
AtSUN3
AtSUN4
HsSUCO
DdSUN2
ScSLP1
ScMPS3
CeSUN1
PfSUN1
AtSUN1
AtSUN2
CeUNC84
HsSUN1
HsSUN2
SpSAD1
HsSUN5
C-terminus
SUN
TM
CC
1
2000

Figure S2.

A.

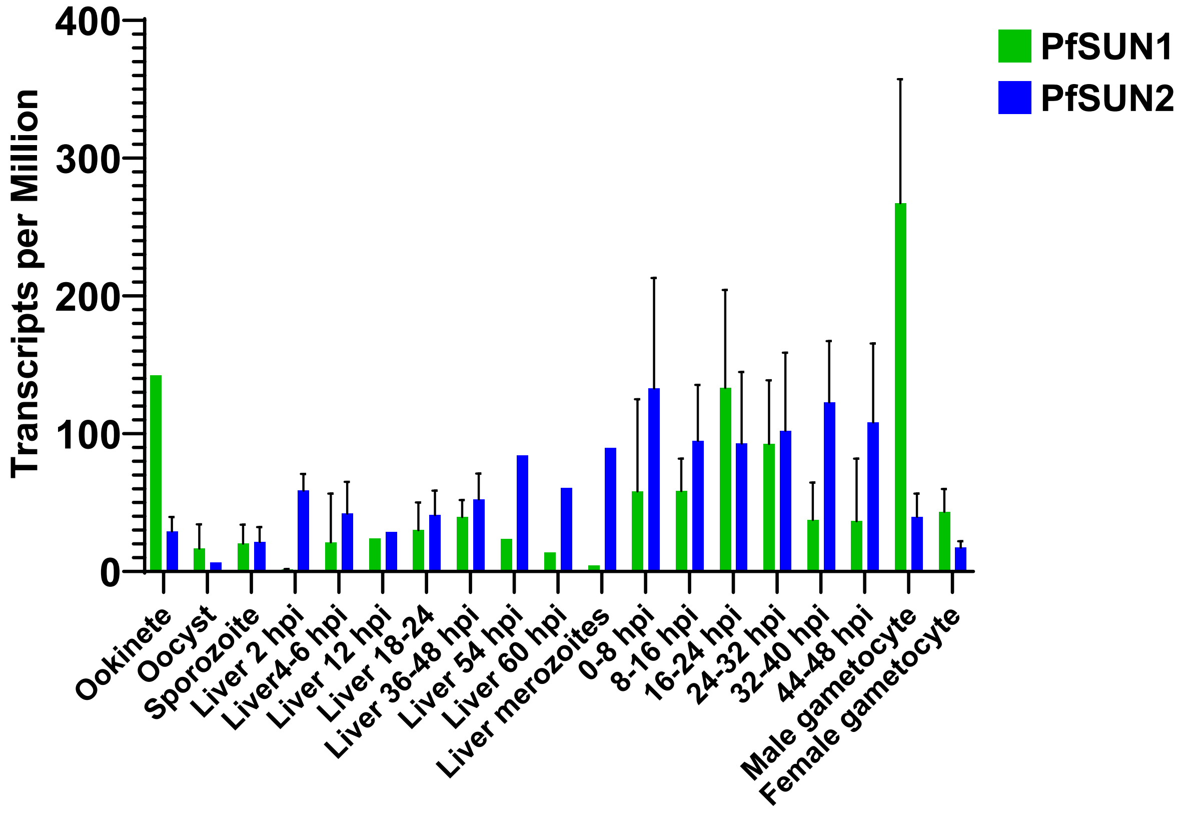

400
300
200
100
0
Transcripts per Million
PfSUN1
PfSUN2
Ookinete
Oocyst
Sporozoite
Liver 2 hpi
Liver4-6 hpi
Liver 12 hpi
Liver 18-24
Liver 36-48 hpi
Liver 54 hpi
Liver 60 hpi
Liver merozoites
0-8 hpi
8-16 hpi
16-24 hpi
24-32 hpi
32-40 hpi
44-48 hpi
Male gametocyte
Female gametocyte


B.

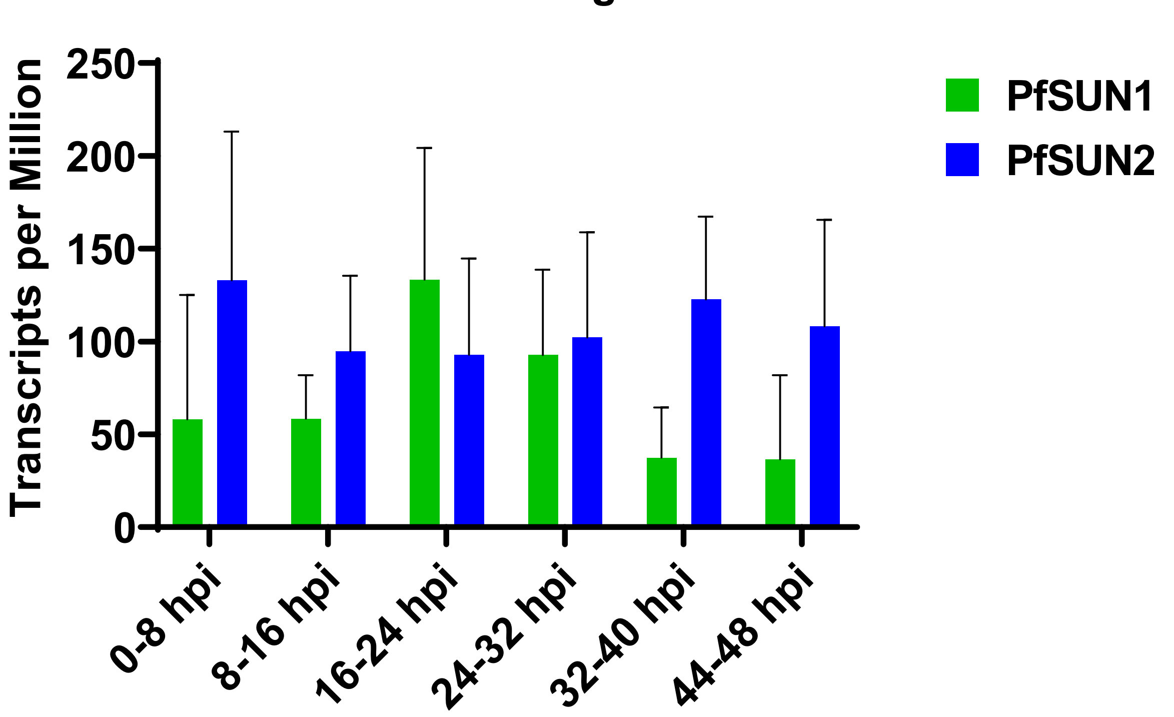

Asexual Stages
250
200
150
100
50
0
Transcripts per Million
PfSUN1
PfSUN2
0-8 hpi
8-16 hpi
16-24 hpi
24-32 hpi
32-40 hpi
44-48 hpi

Figure S3.

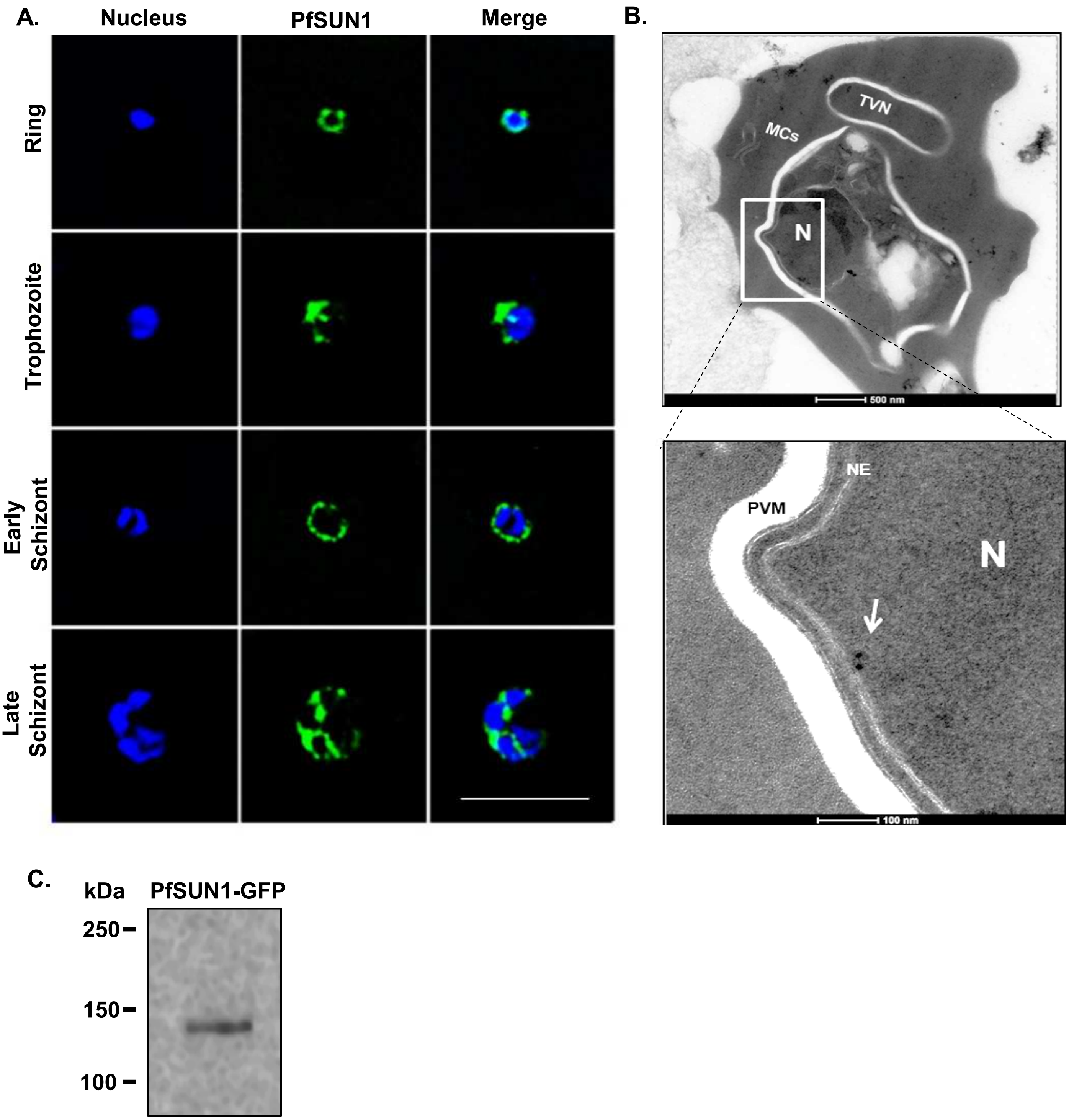
A.
Nucleus
PfSUN1
Merge
Ring
Trophozoite
Early Schizont
Late Schizont
B.
TVN
MCs
N
500 nm
NE
PVM
N
100 nm
C.
kDa
PfSUN1-GFP
250
150
100

**Figure S4.**

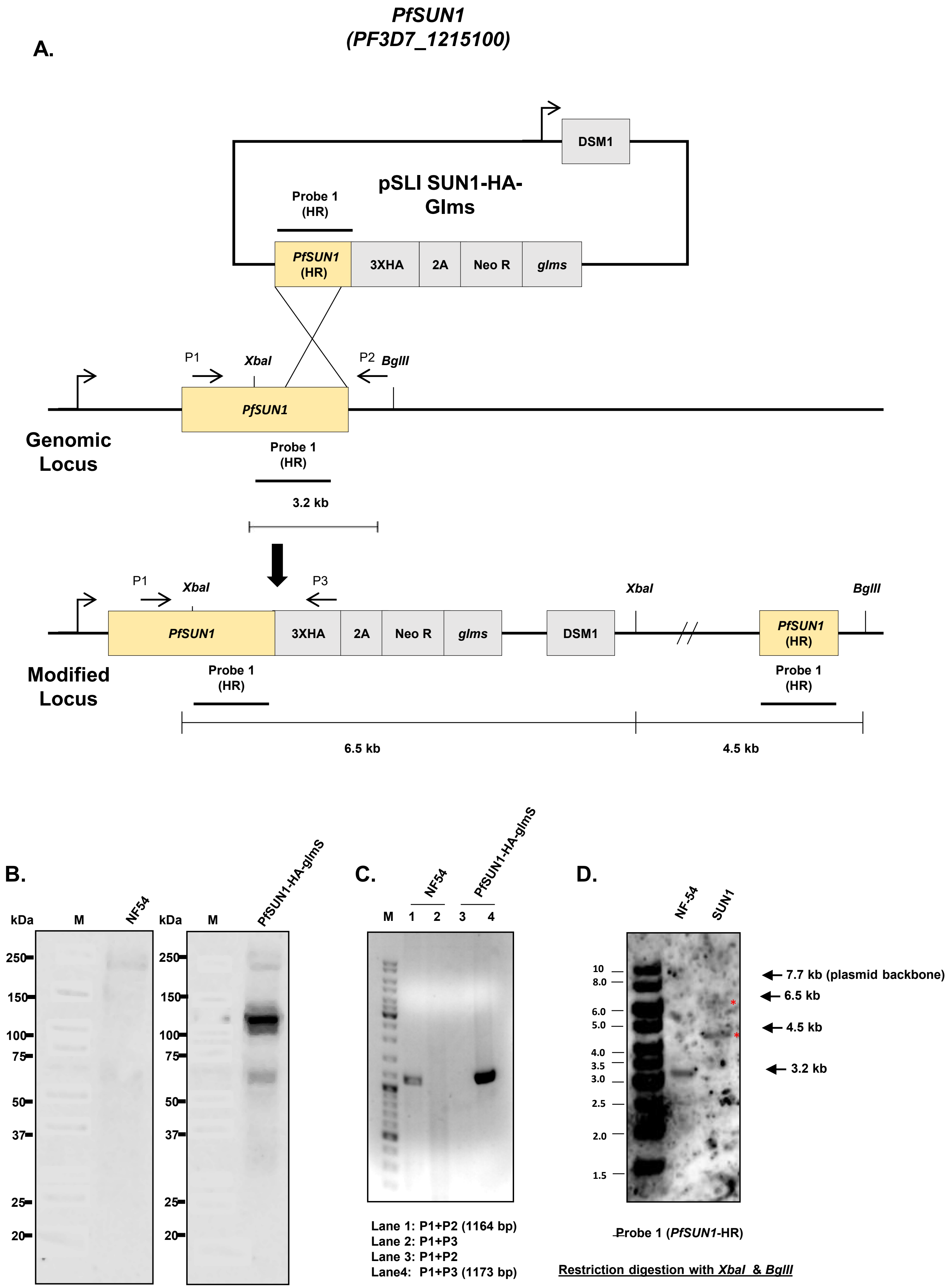

PfSUN1
(PF3D7_1215100)
A.
DSM1
pSLI SUN1-HA-Glms
Probe 1 (HR)
PfSUN1 (HR)
3XHA
2A
Neo R
glms
P1
XbaI
P2
BglII
PfSUN1
Genomic Locus
Probe 1 (HR)
3.2 kb
P1
XbaI
P3
PfSUN1
3XHA
2A
Neo R
glms
DSM1
XbaI
BglII
PfSUN1 (HR)
Modified Locus
Probe 1 (HR)
Probe 1 (HR)
6.5 kb
4.5 kb
B.
kDa
M
NF54
kDa
M
PfSUN1-HA-glmS
250
150
100
75
50
37
25
20
C.
NF54
PfSUN1-HA-glmS
M
1
2
3
4
Lane 1: P1+P2 (1164 bp)
Lane 2: P1+P3
Lane 3: P1+P2
Lane4: P1+P3 (1173 bp)
D.
NF-54
SUN1
10
8.0
6.0
5.0
4.0
3.5
3.0
2.5
2.0
1.5
7.7 kb (plasmid backbone)
6.5 kb
4.5 kb
3.2 kb
Probe 1 (PfSUN1-HR)
Restriction digestion with XbaI & BglII

**Figure S5.**

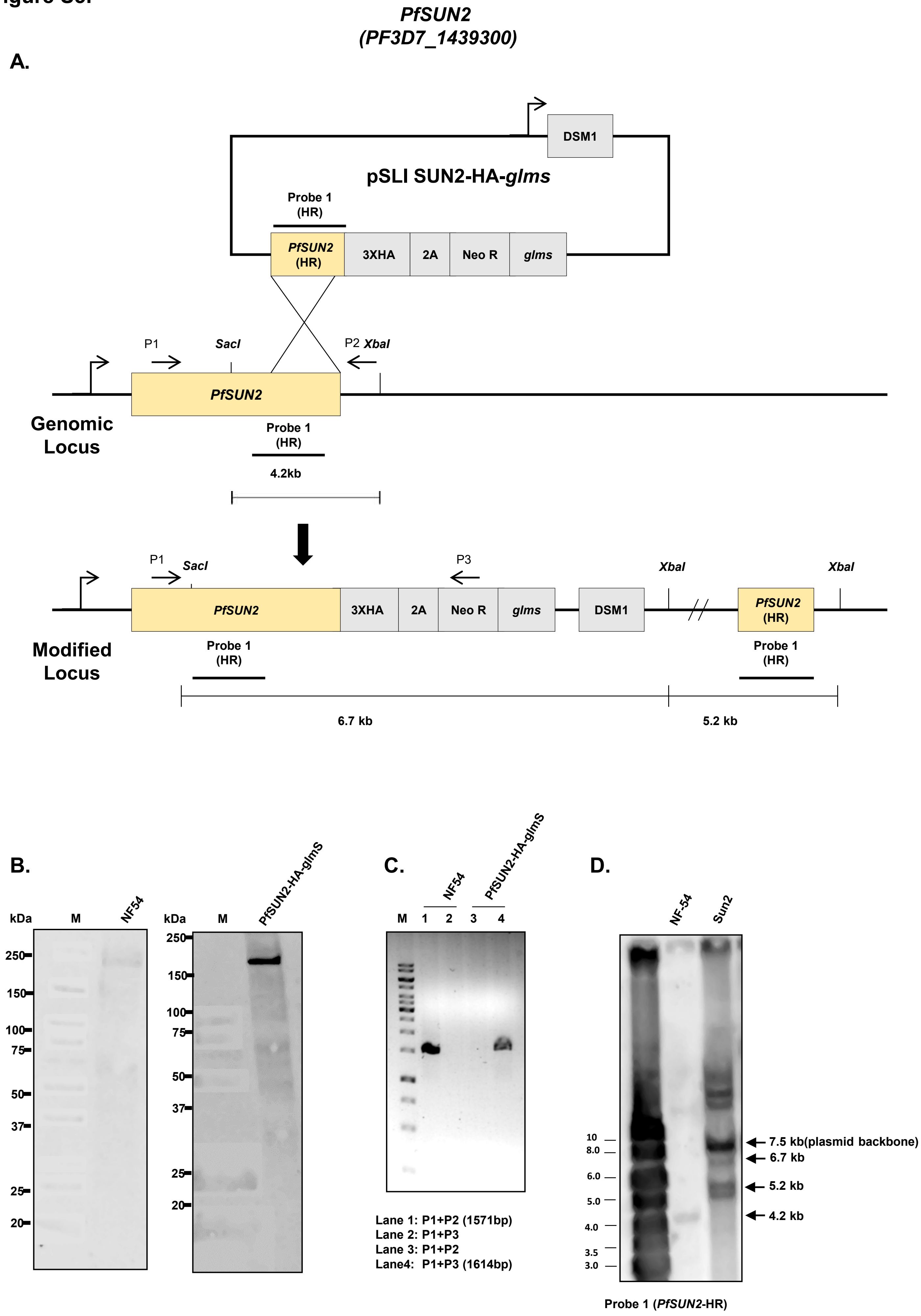

PfSUN2
(PF3D7_1439300)
A.
DSM1
pSLI SUN2-HA-glms
Probe 1 (HR)
PfSUN2 (HR)
3XHA
2A
Neo R
glms
P1
SacI
P2 XbaI
PfSUN2
Genomic Locus
Probe 1 (HR)
4.2kb
P1
SacI
P3
XbaI
XbaI
PfSUN2
3XHA
2A
Neo R
glms
DSM1
PfSUN2 (HR)
Modified Locus
Probe 1 (HR)
Probe 1 (HR)
6.7 kb
5.2 kb
B.
kDa
M
NF54
250
150
100
75
50
37
25
20
kDa
M
PfSUN2-HA-glmS
250
150
100
75
50
37
25
20
C.
NF54
PfSUN2-HA-glmS
M
1
2
3
4
Lane 1: P1+P2 (1571bp)
Lane 2: P1+P3
Lane 3: P1+P2
Lane4: P1+P3 (1614bp)
D.
NF-54
Sun2
10
8.0
6.0
5.0
4.0
3.5
3.0
7.5 kb(plasmid backbone)
6.7 kb
5.2 kb
4.2 kb
Probe 1 (PfSUN2-HR)

**Figure S6.**

A.

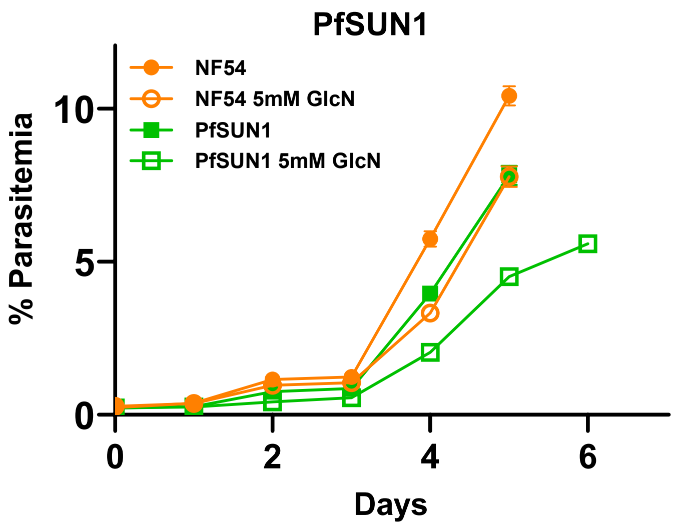

PfSUN1
NF54
NF54 5mM GlcN
PfSUN1
PfSUN1 5mM GlcN
% Parasitemia
10
5
0
0
2
4
6
Days


B.

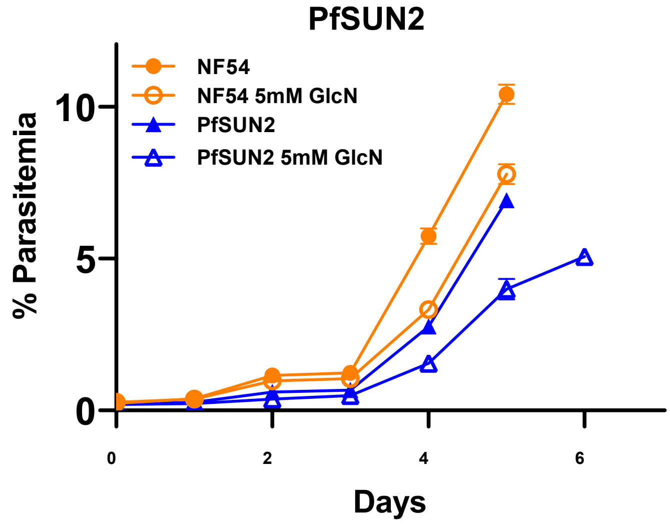

PfSUN2
NF54
NF54 5mM GlcN
PfSUN2
PfSUN2 5mM GlcN
% Parasitemia
10
5
0
0
2
4
6
Days

**Figure S7.**

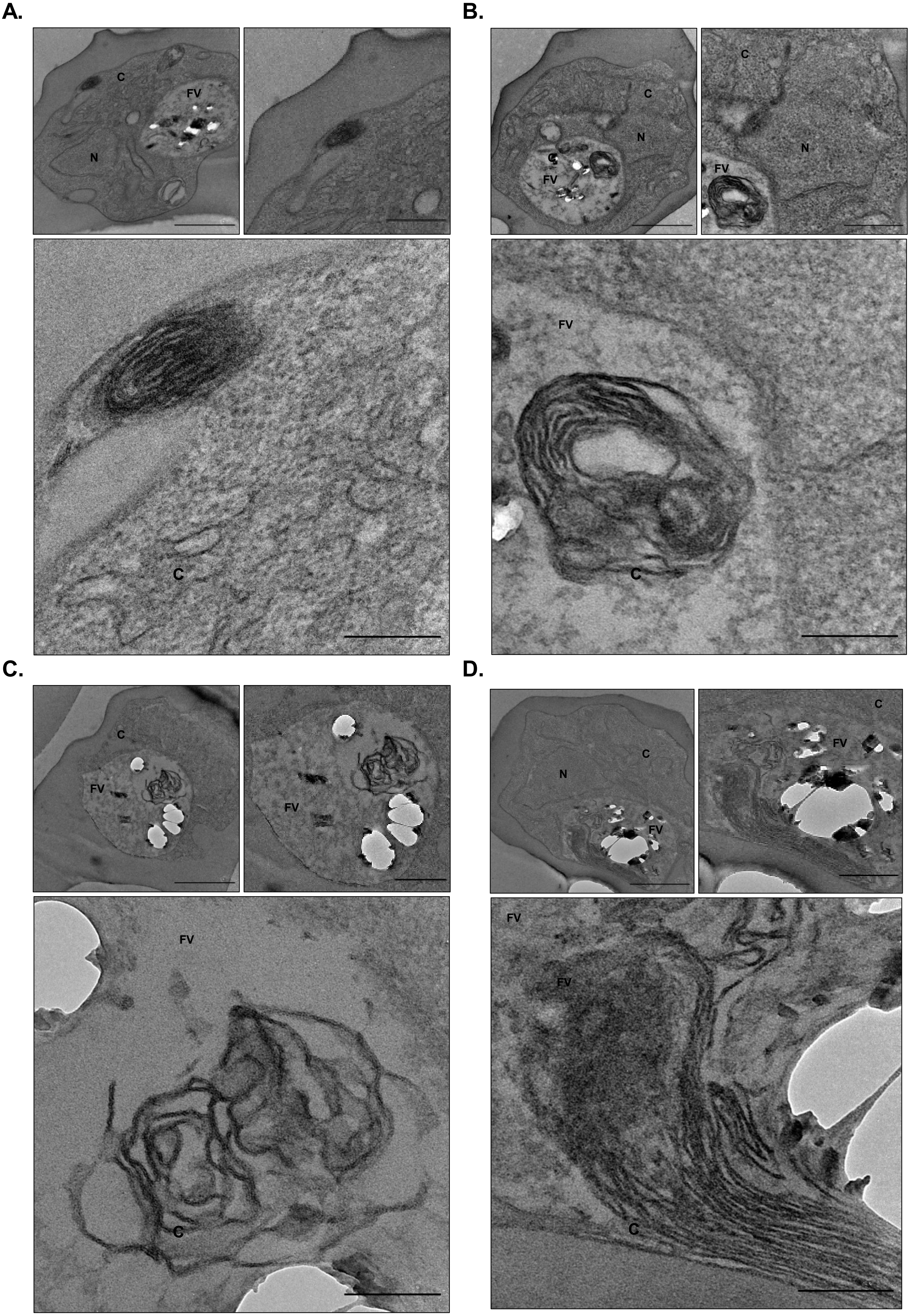

Figure S8.

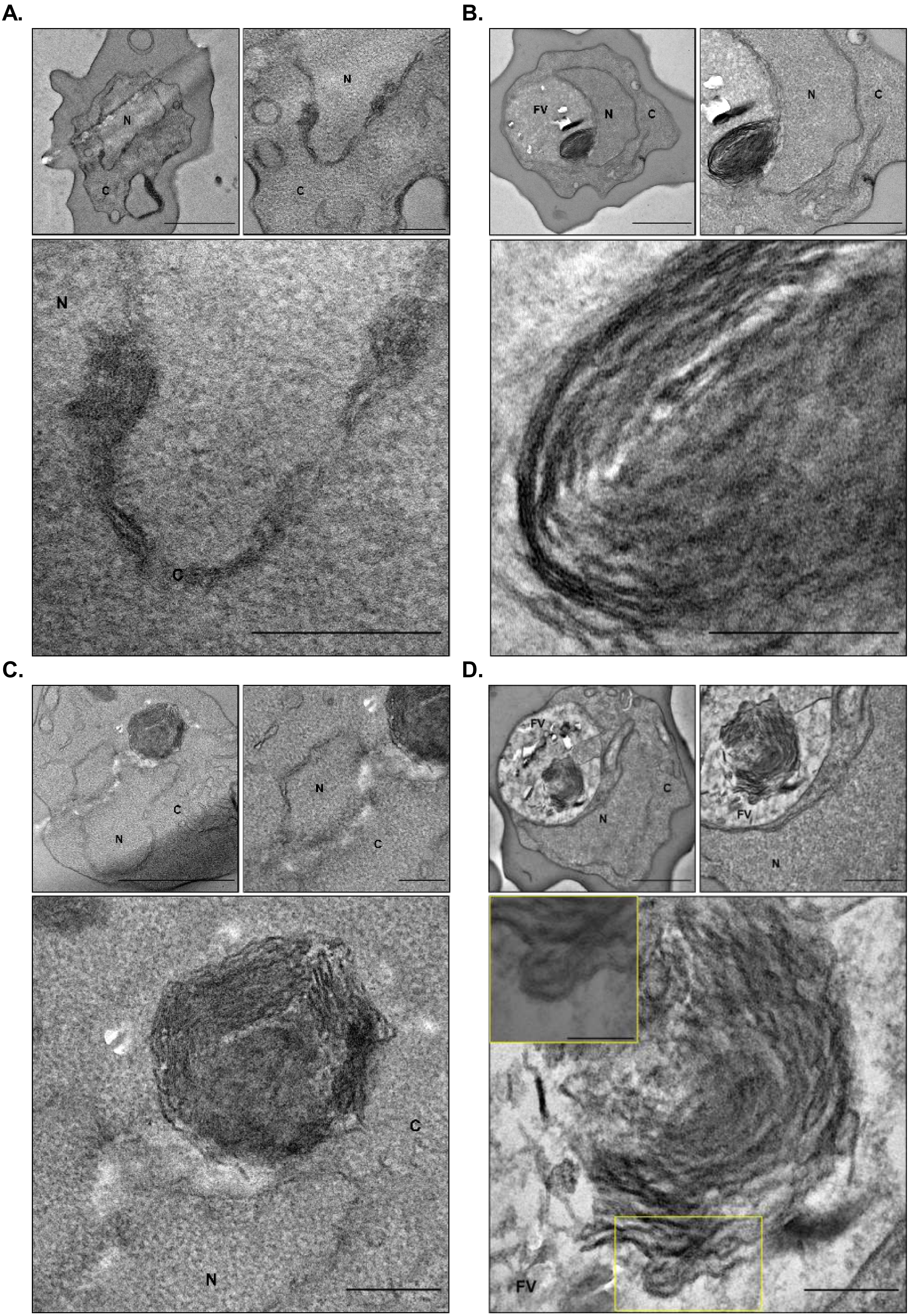

**Figure S9.**

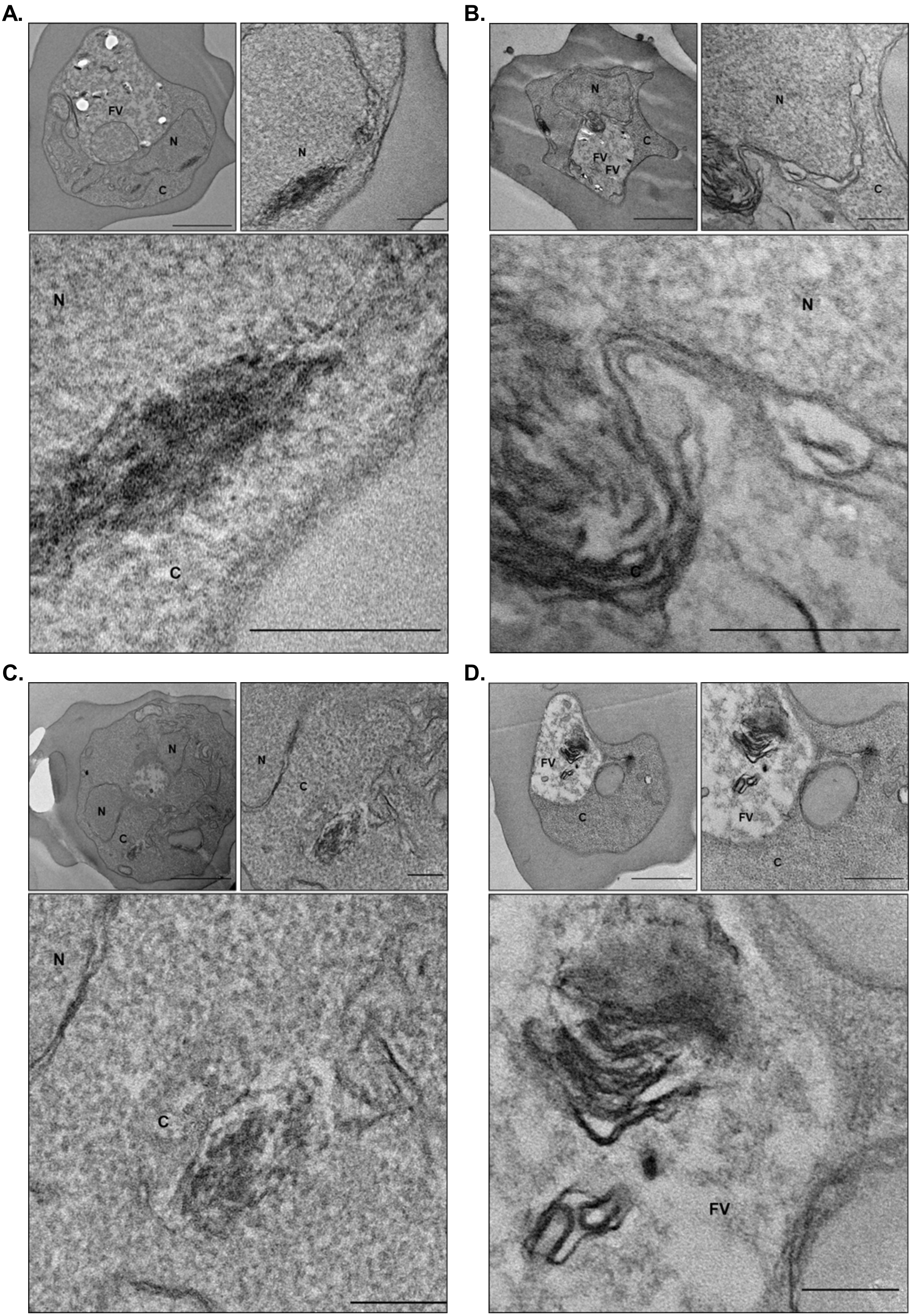

**Figure S10.**

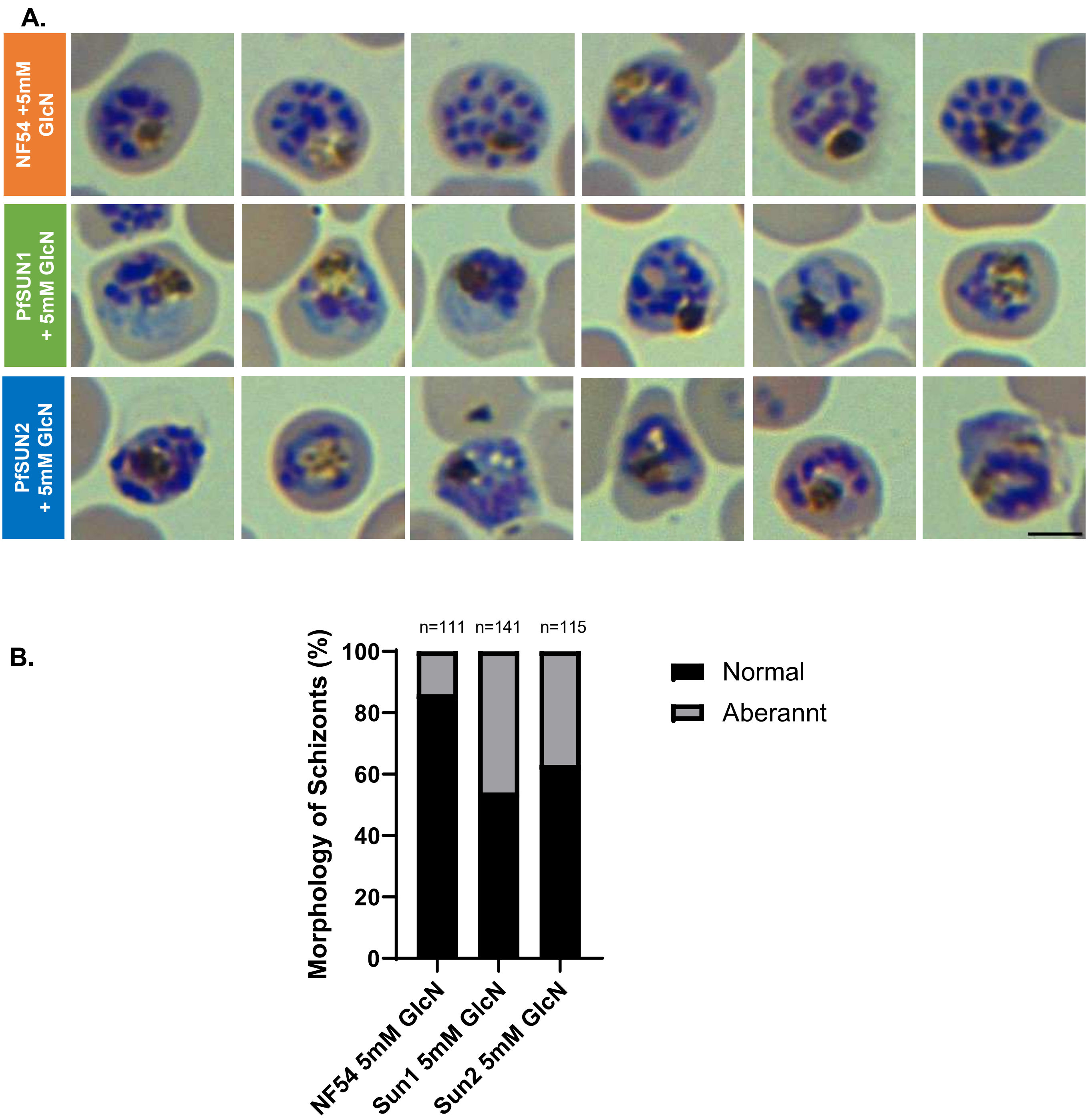
A.
NF54 +5mM GlcN
PfSUN1 + 5mM GlcN
PfSUN2 + 5mM GlcN
B.
n=111
n=141
n=115
Morphology of Schizonts (%)
100
80
60
40
20
0
NF54 5mM GlcN
Sun1 5mM GlcN
Sun2 5mM GlcN
Normal
Aberannt

**Supplementary figures**

**Figure S1. *In silico* analysis of putative SUN domain-containing proteins in *Plasmodium falciparum*. (A and D)** Multiple sequence alignment of amino acid sequences of SUN domain of human (HsSUN2 and HsSUN5), plant (AtSUN5), and putative *P. falciparum* SUN domain proteins named as PfSUN1 **(A)**, and PfSUN2 **(D)**. The alignment was performed using ClustalO (1) with default settings. The secondary structure was fitted onto the alignment using ESPript 3.0 (2). **(B and E)** 3D model structure of the SUN domains of PfSUN1 (522-717, cyan) and PfSUN2 (1-270, cyan), overlaid on the crystal structure of the SUN domain of human SUN2 protein (HsSUN2, PDB ID: 4DXT, magenta). The 3D model structure of the SUN domain of PfSUN1 and PfSUN2 was predicted using I-TASSER (3). PyMol (4) was used for graphical representation. **(C and F)** Sequence comparison of full-length length PfSUN1 **(C)** and PfSUN2 **(F)** among different *Plasmodium* species (50-65% and 50-57% sequence similarity, respectively). Global alignment was performed in Emboss Stretcher (5) under default parameters. **(G)** Evolutionary conservation (phylogenetic analysis) amongst selected SUN domain proteins from different species. Phylogenetic comparison of *P. falciparum* putative SUN domain proteins with 19 SUN domain proteins from different species ranging from plants, yeast, nematode, and human. Multiple sequence alignments of these sequences were obtained from ClustalO and a bootstrapped neighbor-joining tree was obtained. Illustrative visualization of domain architecture of full-length proteins was generated using IBS (Illustrator for biological sequences) (6). Comparative diagrams of SUN-domain proteins depicting protein sizes and domain locations, the positions of transmembrane (TM) (magenta), coiled-coil (CC) (blue), SUN (yellow), are indicated for each protein. Species names and the corresponding UniProt (7) accession numbers for the SUN domain containing proteins: *Homo Sapiens* ( HsSUN1;Q94901 , HsSUN2;Q9UH99, HsSUN5;Q8TC36, HsSUCO;Q94B59), *Plasmodium falciparum* (PfSUN1; Q8I5Q5, PfSUN2;Q8IL76),

*Caenorhabditis elegans* (CeSUN1;Q20924, CeUNC84; Q20745), *Schizosaccharomyces pombe* (SpSAD1;Q09825), *Saccharomyces cerevisiae (ScSLP1;*Q12232, *ScMPS3;*P47069*) Arabidopsis thaliana* (AtSUN1;Q9FF75, AtSUN2;Q9SG79, AtSUN3;F4I316, AtSUN4;F4I8I0, AtSUN5;F4JPE9), *Zea mays* (ZmSUN4;K7VAL9, ZmSUN5; A0A1D6FYJ8) and *Dictyostelium discoideum* (DdSUN2;Q54MI3).

**Figure S2. Compiled expression data of plasmodium SUN domain proteins during Plasmodium life cycle.** Gene expression profiles of **(A)** full life cycle and **(B)** Asexual blood stages were analyzed using RNA-Seq resources implemented in the PlasmoDB database from previously published datasets. All data is for *P. falciparum*, except for liver stages obtained from *P. berghei*. Presented as Mean+SD. (8-19).

**Figure S3. PfSUN1 is localized to the nuclear periphery. A.** IFA imaging of PfSUN1-GFP localization at the nuclear periphery during IDC. Parasite nuclei were stained with DAPI (blue), PfSUN1-GFP was labeled with anti-GFP antibody (green). Scale bar: 10µm. **B.** Transmission electron microscopy using immuno-gold labeling of a PfSUN1-GFP trophozoite detecting PfSUN1 at the inner nuclear membrane. Arrow indicating gold particle. N; Nucleus. NE; Nuclear Envelope. PVM; Parasitophorous Vacuole Membrane. MCs; Maurer's Clefts. TVN; Tubo-vesicular Network, respectively. Scale bar: 500 nm, insert: 100nm. **C.** WB analysis of PfSUN1-GFP parasite lysates using anti-GFP antibody showing that PfSUN1-GFP is translated into a protein of the predicted ~140 KDa.

**Figure S4. An inducible SUN1-HA-*glms* knockdown line was generated using the pSli system.** **(A).** Scheme of the construct used to endogenously tag *Pfsun1*. The predicted structure of the endogenous locus before and after integration is shown. HR, homology region; HA, hemagglutinin epitope tag; 2A, skip peptide; Neo R, neomycin resistance gene: *glmS*, *glms* ribozyme; the scheme includes primers used to detect integration by PCR and restriction of the modified locus by *XbaI* and *BglII***. (B).** Western blot analysis of cellular extracts from NF54 wild type (WT) parasite and PfSUN1-HA-glmS transgenic parasites. Anti-HA antibody detected a clear band at the expected size of PfSUN1-HA (~116kDa) only in PfSUN1-HA-glmS parasites, while no signal was observed in the NF54 WT parasite lysate. **(C).** PCR detection of the transgenic PfSUN1-HA-*glmS* line by amplifying across integration events showing amplicon size difference using primers P1 & P2 (wild type) and P1 & P3 (positive integration). GeneRuler™ DNA Ladder Mix. **(D)** Southern Blot analysis, expected sizes after digestion of gDNA and probing of the blot are noted in the scheme.

**Figure S5. An inducible SUN2-HA-*glms* knockdown line was generated using the pSli system.** **(A)** Scheme of the construct used to endogenously tag *Pfsun2*. The predicted structure of the endogenous locus before and after integration is shown. HR, homology region; HA, hemagglutinin epitope tag; 2A, skip peptide; Neo R, neomycin resistance gene: *glmS*, *glms* ribozyme; the scheme includes primers used to detect integration by PCR and restriction of the modified locus by *XbaI* and *BglII***. (B)** Western blot analysis of cellular extracts from NF54 wild type (WT) parasite and PfSUN2-HA-*glmS* transgenic parasites. Anti-HA antibody detected a clear band at the expected size of PfSUN2-HA (~220kDa) only in PfSUN2-HA-*glmS* parasites, while no signal was observed in NF54 WT parasite lysate. **(C).** PCR detection of the transgenic PfSUN2-HA-*glmS* line by amplifying across integration events showing amplicon size difference using primers P1 & P2 (wild type) and P1 & P3 (positive integration). GeneRuler™ 1kb DNA Ladder. **(D)** Southern Blot analysis, expected sizes after digestion of gDNA and probing of the blot are noted in the scheme.

**Figure S6. Expression of PfSUN1 and PFSUN2 is essential for proper proliferation of blood stage parasites. A.** Growth curve of NF54 (parental line) and PfSUN1-HA-glmS tightly synchronized parasites growing in presence or absence of 5 mM GlcN. Parasitemia was monitored daily using Flow cytometry. Each value is the mean of technical triplicates. Error bars represent Standard Deviation. **B.** Growth curve of NF54 (parental line) and PfSUN2-HA-*glmS* tightly synchronized blood stages parasite grown in presence or absence of 5mM GlcN. Parasitemia was monitored daily using Flow cytometry. Each value is the mean of technical triplicates. Error bars represent Standard Deviation.

**Figure S7. Supplementary micrographs for figure 7A (A-D** corresponds with i-iv respectively**).** Thin section electron micrographs of NF54 parasites grown in the presence of 5mM GlcN. N: Nucleus, C; Cytoplasm, FV; Food Vacuole. Scale bars: Top left ;1µm, top right; 500nm, bottom ;200nm.

**Figure S8. Supplementary micrographs for figure 7A (A-D** corresponds with v-viii respectively**).** Thin section electron micrographs of PfSUN1-HA-*glms* parasites grown in the presence of 5mM GlcN. N: Nucleus, C; Cytoplasm, FV; Food Vacuole. Scale bars: Top left ;1µm, top right; 500nm, bottom ;200nm **(D)** inset; 100nm .

**Figure S9. Supplementary micrographs for figure 7A (E-H** corresponds with ix-xii**).** Thin section electron micrographs of PfSUN2-HA-*glm*s parasites grown in the presence of 5mM GlcN. N: Nucleus, C; Cytoplasm, FV; Food Vacuole. Scale bars: Top left ;1µm, top right; 500nm, bottom ;200nm.

**Figure S10. Knock-down of PfSUN1 and PfSUN2 impairs parasite proliferation during IDC. A.** Representative images of Giemsa-stained tightly synchronized schizonts of each line grown in the presence of 5mM GlcN and used for EM presented in figure 5E. **B.** bar graph quantifies the proportion of normal and aberrant schizonts for each parasite line. Scale bar: 5 µm.


1. Sievers F & Higgins DG (2014) Clustal Omega, accurate alignment of very large numbers of sequences. *Multiple sequence alignment methods*, (Springer), pp 105-116.
2. Gouet P, Robert X, & Courcelle E (2003) ESPript/ENDscript: Extracting and rendering sequence and 3D information from atomic structures of proteins. *Nucleic Acids Res* 31(13):3320-3323.
3. Yang J*, et al.* (2015) The I-TASSER Suite: protein structure and function prediction. *Nat Methods* 12(1):7-8.
4. Schrödinger L & DeLano W (2020) PyMOL. *The PyMOL Molecular Graphics System, Version* 2.
5. Rice P, Longden I, & Bleasby A (2000) EMBOSS: the European Molecular Biology Open Software Suite. *Trends Genet* 16(6):276-277.
6. Liu W*, et al.* (2015) IBS: an illustrator for the presentation and visualization of biological sequences. *Bioinformatics* 31(20):3359-3361.
7. UniProt C (2015) UniProt: a hub for protein information. *Nucleic Acids Res* 43(Database issue):D204-212.
8. Toro-Moreno M, Sylvester K, Srivastava T, Posfai D, & Derbyshire ER (2020) RNA-Seq Analysis Illuminates the Early Stages of Plasmodium Liver Infection. *mBio* 11(1).
9. Chappell L*, et al.* (2020) Refining the transcriptome of the human malaria parasite Plasmodium falciparum using amplification-free RNA-seq. *BMC Genomics* 21(1):395.
10. Lindner SE*, et al.* (2019) Transcriptomics and proteomics reveal two waves of translational repression during the maturation of malaria parasite sporozoites. *Nat Commun* 10(1):4964.
11. Caldelari R*, et al.* (2019) Transcriptome analysis of Plasmodium berghei during exo-erythrocytic development. *Malar J* 18(1):330.
12. Toenhake CG*, et al.* (2018) Chromatin Accessibility-Based Characterization of the Gene Regulatory Network Underlying Plasmodium falciparum Blood-Stage Development. *Cell Host Microbe* 23(4):557-569 e559.

13. Gomez-Diaz E*, et al.* (2017) Epigenetic regulation of Plasmodium falciparum clonally variant gene expression during development in Anopheles gambiae. *Scientific reports* 7:40655.
14. Zanghi G*, et al.* (2018) A Specific PfEMP1 Is Expressed in P. falciparum Sporozoites and Plays a Role in Hepatocyte Infection. *Cell reports* 22(11):2951-2963.
15. Lopez-Barragan MJ*, et al.* (2011) Directional gene expression and antisense transcripts in sexual and asexual stages of Plasmodium falciparum. *BMC Genomics* 12:587.
16. Bartfai R*, et al.* (2010) H2A.Z demarcates intergenic regions of the plasmodium falciparum epigenome that are dynamically marked by H3K9ac and H3K4me3. *PLoS Pathog* 6(12):e1001223.
17. Wichers JS*, et al.* (2019) Dissecting the Gene Expression, Localization, Membrane Topology, and Function of the Plasmodium falciparum STEVOR Protein Family. *mBio* 10(4).
18. Otto TD*, et al.* (2010) New insights into the blood-stage transcriptome of Plasmodium falciparum using RNA-Seq. *Mol Microbiol* 76(1):12-24.
19. Lasonder E*, et al.* (2016) Integrated transcriptomic and proteomic analyses of P. falciparum gametocytes: molecular insight into sex-specific processes and translational repression. *Nucleic Acids Res* 44(13):6087-6101.
